# A genome-wide genetic screen uncovers novel determinants of human pigmentation

**DOI:** 10.1101/2021.09.29.462413

**Authors:** Vivek K. Bajpai, Tomek Swigut, Jaaved Mohammed, Josh Tycko, Sahin Naqvi, Martin Arreola, Tayne C. Kim, Neha Arora, Jonathan K. Pritchard, Michael C. Bassik, Joanna Wysocka

## Abstract

The skin color is one of the most diverse human traits and is determined by the quantity, type and distribution of melanin. Here, we leverage light scattering properties of melanin to conduct a genome-wide CRISPR-Cas9 screen for novel regulators of melanogenesis. We identify functionally diverse genes converging on melanosome biogenesis, endosomal transport and transcriptional/posttranscriptional gene regulation, most of which represent novel associations with pigmentation. A survey of transcriptomes from diversely pigmented individuals reveals that the majority of genes discovered in our screen are upregulated in dark skin melanocytes, in agreement with their melanin-promoting function and potential contribution to skin color variation. This association is further buttressed by the significant skin color heritability enrichment in the vicinity of these genes. Taken together, our study presents a novel approach to assay pigmentation and uncovers a plethora of melanogenesis regulators, with broad implications for human variation, cell biology and medicine.

**One Sentence Summary:** Genetic screen uncovers genes involved in human melanogenesis, many of which are differentially expressed in individuals of diverse skin color.

## Main text

The skin and hair color are genetically derived traits that are highly variable between and within human populations and determined by the quantity, type and distribution of melanin (*1*). Melanocytes (MC), which developmentally originate from the neural crest and reside in the epidermis, synthesize melanin within subcellular organelles called melanosomes, which upon maturation are transported to the surrounding epidermal keratinocytes resulting in skin and hair pigmentation (**Fig. 1A**) (*1*). Key insights into human pigmentation have come from mapping genes involved in hypo- and hyper-pigmentation diseases and from candidate and genome-wide association studies (GWAS) of normal-range skin and hair color variation in human populations (*1, 2*). Furthermore, work in model organisms, in particular studies of mouse coat color, revealed genes and pathways involved in pigmentation, many of which converge on the melanin synthesis pathway (*3, 4*). Nonetheless, current genetic knowledge can explain only a small fraction of skin color variation in humans suggesting that many genes controlling pigmentation in our species remain undiscovered (*2*).

**Fig. 1.**
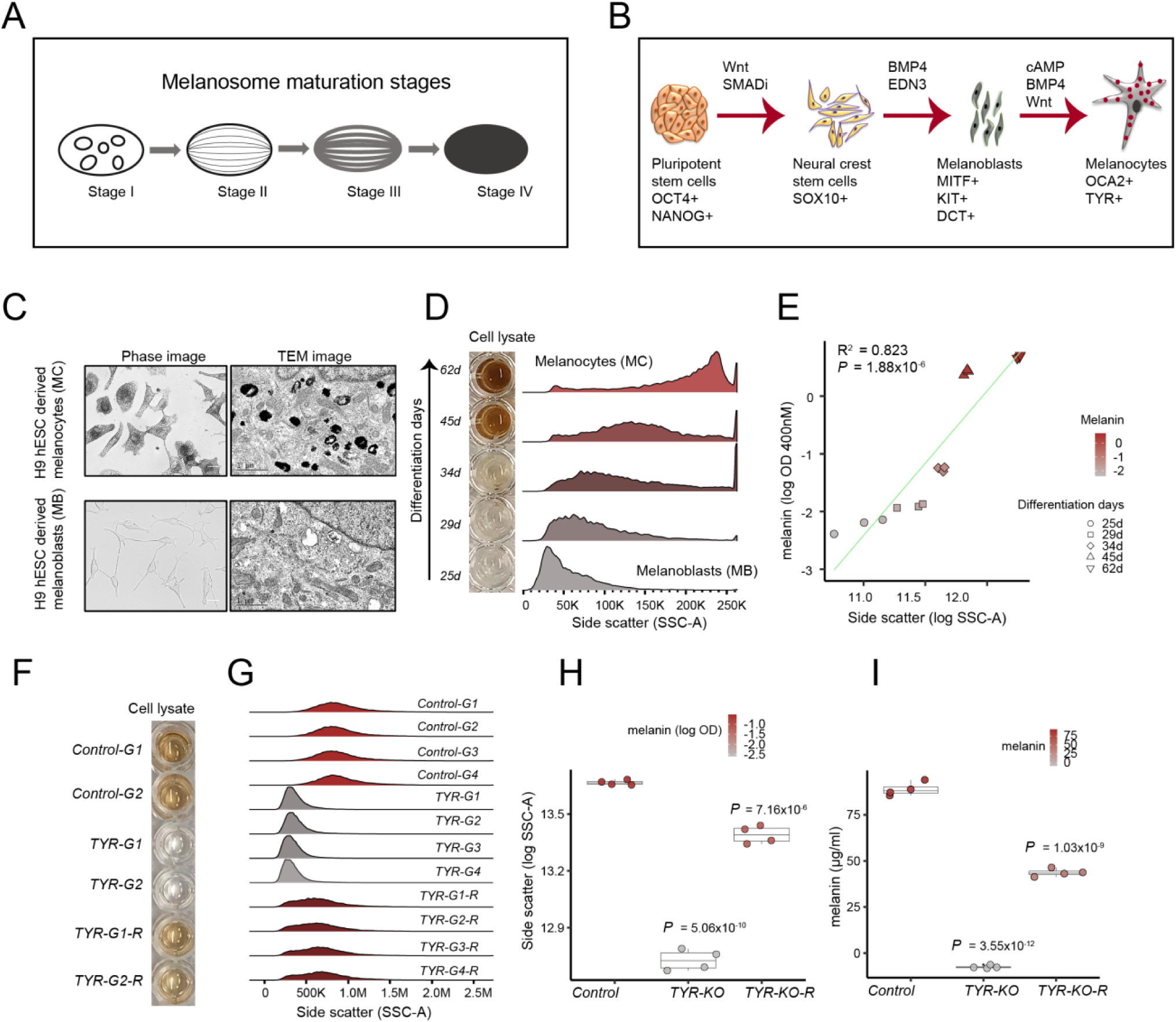
Melanin concentration determines light scattering property of pigment cells. (**A)** Schematic of four stages of melanosome maturation. Stage I melanosomes contain intraluminal vesicles, stage II melanosomes deposit parallel PMEL amyloid fibrils on which melanogenesis takes place. Stage III and IV melanosomes correspond to partially and fully melanized states, respectively. (**B)** Schematic of the differentiation process. PSC were differentiated into neural crest cell, melanoblasts and melanocytes in a sequential manner, following a previously established protocol (*7*). The key growth factors and signaling molecules driving differentiation are shown above the arrows and lineage markers characterizing each stage are shown below each cell type. (**C)** Differentiation of H9 hESC derived melanoblasts (MB, bottom panels) into melanocytes (MC, top panels) results in increased melanized stage III-IV melanosomes as shown by phase and transmission electron microscopy (TEM) (n=3 independent differentiations). (**D)** Increased melanin concentration during melanocyte differentiation from hESC (days of differentiation are indicated on the left) is accompanied by an increase in light scattering as measured by side scatter (SSC) parameter of flow cytometer. (**E**) Relationship between melanin (OD 400nM) and SSC as modeled by linear regression. R^2^ = 0.823, *P* = 1.88×10^−6^. (**F**) CRISPR-Cas9 mediated loss of *TYR* (tyrosinase) gene makes pigmented MNT1 melanoma cells amelanotic and re-expression of *TYR* transgene recovers the pigmentation (n = 4 sgRNAs treatments). Cell lysates from two representative knockout and rescue experiments are shown. (**G)** *TYR* loss significantly reduces the side scatter and *TYR* re-expression restores it. Four different sgRNA against TYR and control sgRNAs are labeled as G1, G2, G3 and G4 after the gene name. **(H)** Boxplot showing side scatter (log SSC-A) changes in response to *TYR* loss and rescue in comparison to control-edited cells. Filled color within dots corresponds to total melanin (log OD at 400nM). Box plots show median and IQR, whiskers are 1.5 × IQR. Significance tested for control, TYR-KO and TYR-KO-R groups using ANOVA followed by two-sided Welch t-Test with Benjamini & Hochberg correction. *P* values shown are relative to control. (**I**) Total melanin concentration calculated for loss and rescue of *TYR* (tyrosinase) experiment (n=4 sgRNAs treatment). Melanin content was determined spectrophotometrically by measuring OD (at 400nM) of cell lysates (6×10^5^ cells in 120 µl 1M NaOH with 10% DMSO) and comparing against a standard curve generated using synthetic melanin as shown in (S1D). Box plots showing median melanin concentration (µg/ml) and IQR, whiskers are 1.5 × IQR. Significance was tested for four sgRNA and TYR rescue treatment with ANOVA followed by two-sided Welch t-Test with Benjamini & Hochberg correction. *P* values shown are relative to control. Scale bars: A, phase images = 25µM.

Melanin is a general term for a heterogenous and structurally ill-defined biopolymer (*5*) that encompasses two forms, namely black/brown eumelanin and red/yellow pheomelanin. The quantity and type of produced melanin determines its physicochemical properties such as high refractive index, which forms the basis of melanin’s characteristic light absorbance and scattering properties (*5*), thought to be critical for protecting the skin from the sun exposure-related damage. Light scattering measured as a side scatter (a.k.a. 90° scatter) parameter in flow cytometry reflects internal complexity of the cells (*6*). We investigated the extent to which changes in cellular melanin content influence light scattering measured as side scatter (SSC) and whether SSC can be used to capture dynamic changes in the melanin levels upon inducing genetic perturbations. Here, we demonstrate that cellular melanin concentration indeed determines light scattering (SSC) properties of pigment cells. Taking advantage of this unique feature and using SSC as a proxy for melanin levels, we performed a CRISPR-Cas9 based genetic screen to identify novel regulators of human pigmentation.

To establish a link between melanin content and SSC, we employed two different cellular models of human melanogenesis. First, we investigated the relationship between SSC and pigmentation in a developmental context, where we modeled melanogenesis using pluripotent stem cells (**Fig. 1B**) (*7*). To this end, we differentiated *SOX10::GFP* reporter H9 human embryonic stem cells (hESC) to SOX10+ neural crest (NC) cells, then coaxed NC towards unpigmented precursors of MC called melanoblasts (MB), and finally differentiated those into mature pigmented MC, confirming differentiation with appropriate lineage markers at each step (**Fig. S1A, B**). The *in vitro* derived MC had high melanin content, as determined by optical density (OD) measurements of cell lysate at 400nM wavelength and comparing against a standard curve generated using synthetic melanin (*8*) (**Fig. S1C, D**). During maturation, they gradually accumulated pigmented melanosomes, as determined by phase and transmission electron microscopy (TEM) imaging (**Fig. 1C**). Furthermore, RNA-seq expression analysis of MB and MC populations derived from two different pluripotent stem cell (PSC) lines revealed strong upregulation of melanogenic genes (e.g. *OCA2, TYR, SLC45A2, OA1, PMEL*) during maturation, as expected (**Fig. S1E**). After confirming the identity of MB and MC, we measured the total melanin content and SSC of MB as they matured into MC (**Fig. S1F**). We observed that an increase in melanin content, as measured by OD at 400nM, was accompanied by an increase in SSC (**Fig. 1D, E**) and there was a strong linear correlation between SSC and melanin content (R^2^ = 0.82, *P* = 1.88×10^−6^) (**Fig. 1E**).

In an orthogonal approach to establish the relationship between SSC and pigmentation, we used MNT1 melanoma cell line, which is transcriptionally similar to normal primary human MC (*9*), and has been previously used as a model to study melanogenesis (*10-13*). Using several independent RNA guides (G1-G4), we deleted the key melanogenic gene tyrosinase (*TYR*) in MNT1 cells, which resulted in loss of pigmentation, and decreased melanin absorbance measurements (**Fig. 1F, I** and **Fig. S1G**). Importantly, loss of melanin was accompanied by a significant decrease in SSC (**Fig. 1G, H**). TEM images of *TYR* mutant cells showed preponderance of stage II unpigmented melanosomes and lack of stage III/IV melanized melanosomes, thus providing evidence that melanosome maturation state is directly linked to changes in SSC (**Fig. S1G**). Furthermore, overexpression of wild type copy of *TYR* restored both pigmentation and SSC (**Fig. 1F, G, H, I**). Together, these experiments establish that perturbations in melanosome biogenesis of pigment cells can be monitored by measuring SSC.

We reasoned that the close correlation between melanin content and SSC could be used as a paradigm for a CRISPR-Cas9 genetic screen, in which loss of genes important for melanosome biogenesis and maturation would result in diminished SSC (**Fig. 2A**). We generated a clonal MNT1 cell line inducibly expressing SpCas9 nuclease upon doxycycline treatment (Cas9-MNT1 cell line; **Fig. S2A**), and infected it with a genome-wide lentiviral sgRNA library (with 10 targeting-guides per gene) (*14*), such that each cell expressed a single sgRNA. In addition, the library also included negative control sgRNAs, such as non-targeting guides (i.e. no binding sites in the genome) and safe-targeting control guides targeting genomic loci with no annotated function (*14*). Following puromycin selection to kill noninfected cells, expression of Cas9 was induced to carry out gene-editing, and after two weeks, cells from the top and bottom 10% of the SSC distribution were FACS sorted (**Fig. 2A, B**). As expected, the low SSC fraction was enriched for hypo-pigmented cells relative to the high SSC fraction (**Fig. 2B**). Genomic DNA was subsequently isolated and the frequencies of sgRNAs in both populations (low and high SSC) were measured by deep sequencing and analyzed using the Cas9 high-Throughput maximum Likelihood Estimator (CasTLE) algorithm, which provides a confidence score for the effect of each gene (*15*). Analysis of CasTLE scores in two independent biological replicates (i.e. independent sgRNA infections and FACS sorting post Cas9 induction) revealed good concordance (R^2^ = 0.59) between the replicates (**Fig. S2B**). We focused our subsequent analysis on genes with a positive CasTLE effect (namely those whose loss corresponded to a reduction in SSC), and which are therefore predicted to have a melanin-promoting function. Although our screen also identified genes with a negative CasTLE effect (e.g. those enriched in the high SSC sorted fraction) we reasoned that gain of SSC may be more challenging to interpret, as it can be affected by other factors which increase cellular granularity (*16*).

**Fig. 2.**
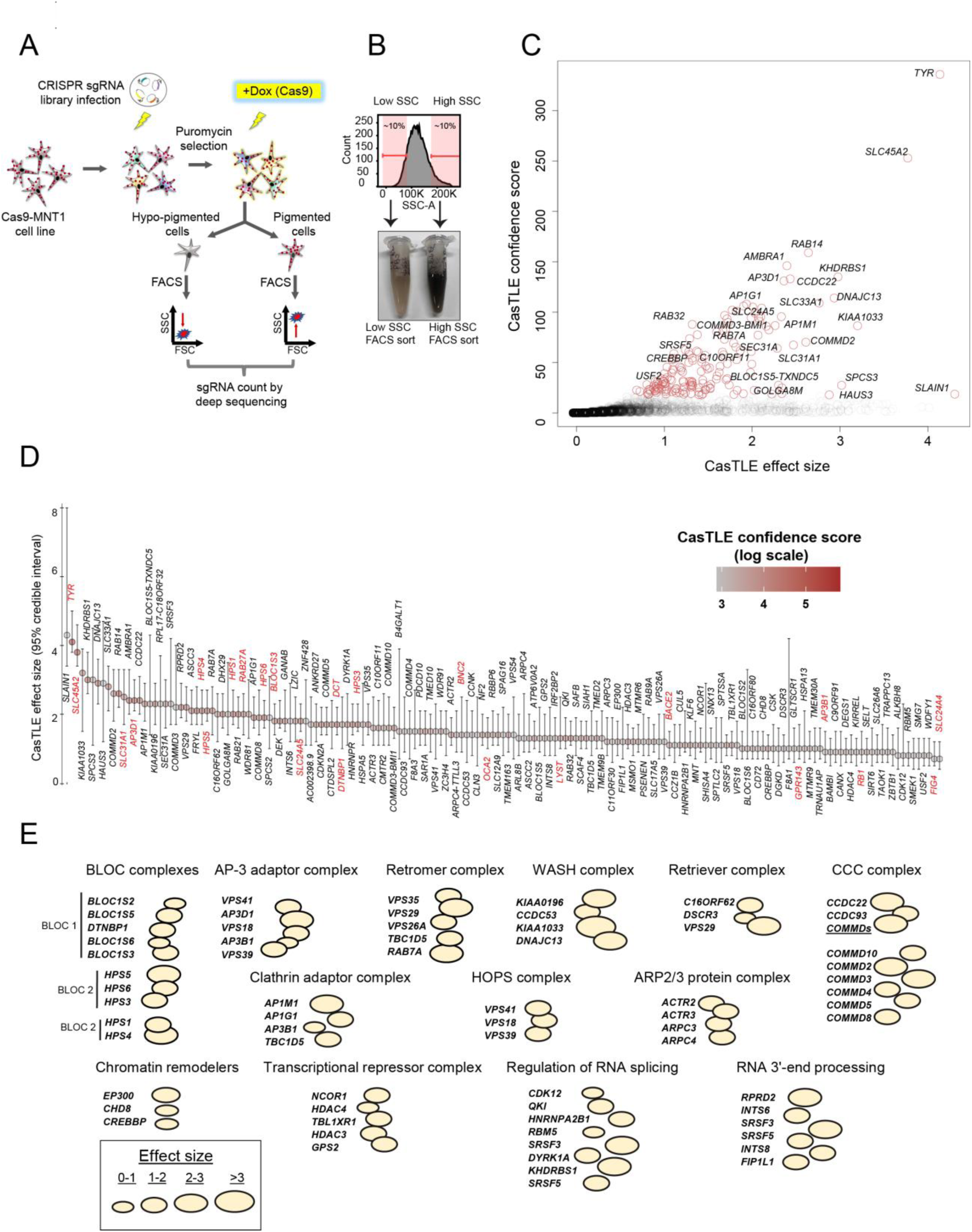
Genome-wide CRISPR-Cas9 screen for regulators of human melanogenesis. (**A)** Schematic of the screen design. (**B)** FACS cell sorting on low and high side scatter (SSC) enriches for hypopigmented and hyperpigmented cells, respectively. (**C)** CasTLE likelihood ratio test (*15*) analysis of two independent genome-wide pigmentation screens. Genes at 10% FDR cutoff are shown as brown circles. (**D)** The maximum effect size (center value) estimated by CasTLE from two independent genome-wide pigmentation screens with ten independent sgRNAs per gene. Bars, 95% confidence interval. Melanogenesis regulators as sorted by CasTLE effect size and filled color represent CasTLE confidence score (log scale). Previously known pigmentation genes are highlighted in red. (**E**) Classification of screen hits based on biological function and presence in common macromolecular complexes. Bubble size indicates CasTLE effect size.

Our screen identified 169 putative melanin-promoting genes (i.e. genes whose deletion was associated with diminished SSC) in common between the two biological replicates at 10% false discovery rate (FDR) (**Fig. 2C, D** and **Table S1**). Importantly, 23 of the 169 screen hits correspond to known melanogenic genes (e.g. *TYR, OCA2, SLC45A2, DCT, SLC24A5, HPS1, HPS3-6, LYST, AP3D1* and others, highlighted in red in **Fig. 2D**), whereas the majority represent novel candidates for regulation of melanogenesis. At lower statistical confidence, between 10-20% FDR, the screen discovered additional 75 hits (**Table S1**), which included several known pigmentation genes (e.g. *MYO5A, ATP7A* and *RAB1A* etc.) suggesting that the 10% FDR cutoff we selected for our analysis is conservative and may miss additional genuine hits. We also note that our screening paradigm is not expected to identify genes that regulate pigmentation indirectly, such as those associated with melanocyte development, survival and migration, but which nonetheless may influence pigmentation *in vivo*.

Gene ontology analysis of screen hits revealed enrichment for melanosome organization, pigment granule organization, vesicle organization and intracellular transport (**Fig. S3A** and **Table S2**). Interestingly, the screen hits were not only enriched for previously known melanosome biogenesis/maturation related factors such as BLOC (biogenesis of lysosomal organelles complex) and AP-3 adaptor complexes (*17*), but also encompassed genes whose products form molecular complexes implicated in endosomal recycling (such as retromer, retriever, WASH, ARP2/3, CCC (COMMD-CCDC22-CCDC93) complexes) (*18*) (**Fig. 2E, S3B** and **Table S2**). These results suggest hitherto underappreciated role of endosome recycling pathways in melanosome biogenesis and human pigmentation. Hits also included genes regulating chromatin modification and remodeling, as well as RNA splicing and processing, suggesting that both transcriptional and post-transcriptional modes of gene regulation contribute towards melanogenesis (**Fig. 2E**).

To validate select hits representing a range of biological functions and screen effect sizes, we introduced sgRNAs against *AP1G1, CCDC22, COMMD3, KHDRBS1, KLF6, SLC12A9, SLC33A1, TYR, KIAA1033 (WASHC4)* and *WDR81* into the Cas9-MNT1 cell line (**Fig. S4A** and **Fig. 3A, B**). Multiple knockout experiments using three distinct sgRNAs against each of the aforementioned genes confirmed their regulatory role in human melanogenesis, as knocking out each gene significantly reduced pigmentation, albeit to varying degrees (**Fig. 3A, B**). As expected, the reduced pigmentation phenotype was accompanied by a diminished SSC (**Fig. S4B**). Notably, our validation experiments were done on bulk cell populations, which likely contain a mixture of heterozygous, homozygous and unedited cells. Thus, melanin measurements underestimate the loss of gene pigmentation phenotype expected from a complete (homozygous) loss of gene function.

**Fig. 3.**
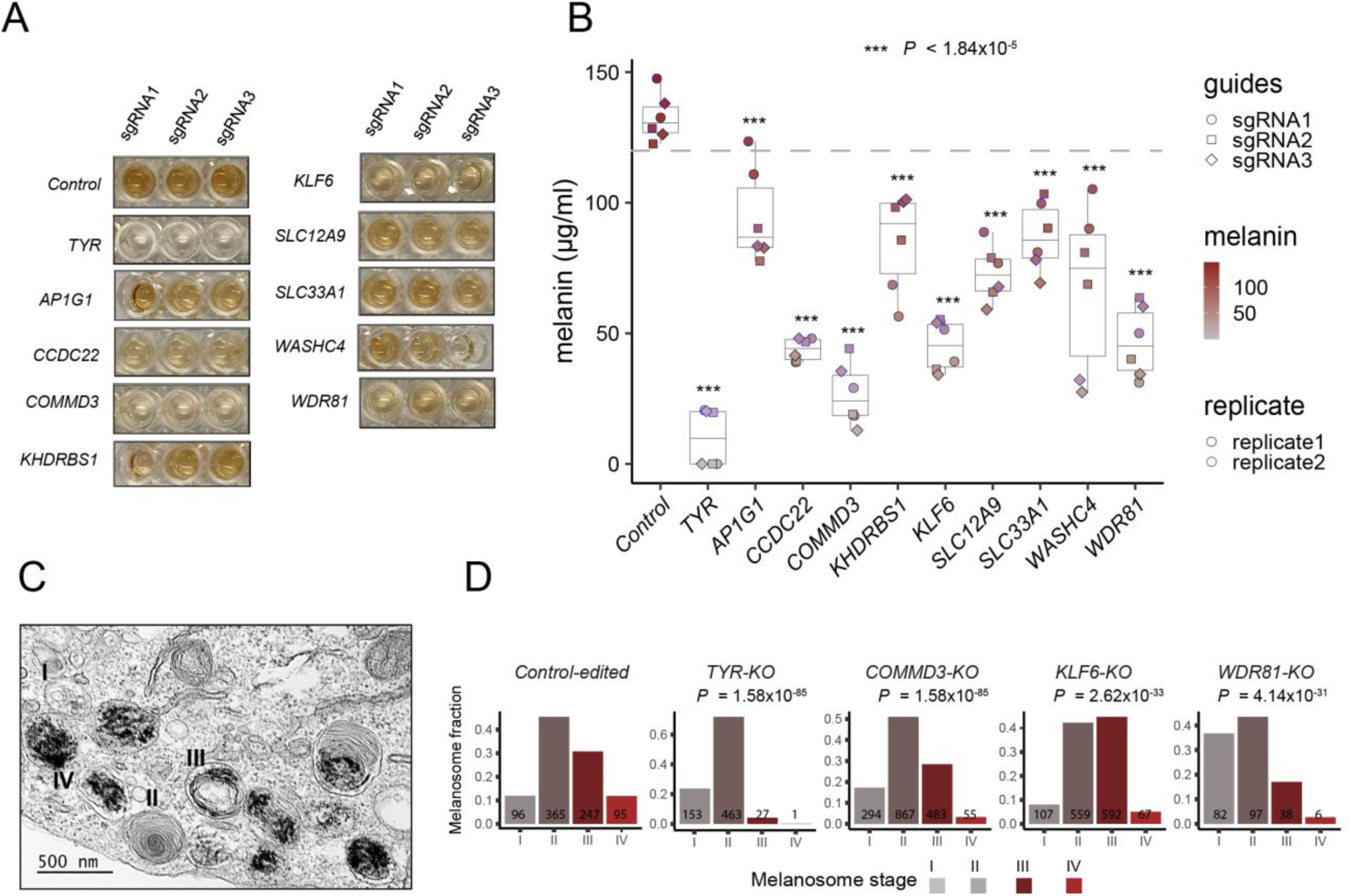
Novel pigmentation genes affect different aspects of melanosome biogenesis and maturation. (**A**) CRISPR-Cas9 based validation of select novel pigmentation screen hits along with negative (safe-targeting sgRNAs) and positive (*TYR*) controls. Cas9-MNT1 cell lysates showing gross changes in melanin levels upon knockout of indicated genes using three different sgRNAs. (**B)** Total melanin content was determined spectrophotometrically by measuring OD (at 400nM) of cell lysates (6.0×10^5^ cells in 120 µl 1M NaOH with 10% DMSO) and comparing against a standard curve generated using synthetic melanin. Melanin measurements were done at two different time points (labeled as replicates) for three sgRNAs against each gene including safe-targeting control sgRNAs. Box plots show median melanin concentration (µg per ml) and IQR, whiskers are 1.5 × IQR. Significance tested with ANOVA followed by two-sided pairwise Welch t-test with Benjamini & Hochberg correction. *P* values relative to control-edited cells ranged from smallest 2.46×10^−20^ (for *TYR*-KO cells) to largest 1.84×10^−5^ (for *AP1G1*-KO cells). *** indicate that for all groups *P* < 1.84×10^−5^. (**C)** Representative TEM image showing melanosomes at stages I to IV of maturation. (**D)** Quantification of melanosomes at different stages of maturation among different indicated gene knockouts including control guides. Bar represents fraction of melanosome counts in stages I-IV. Total counts are displayed on each bar. Overall differences in distribution of melanosome fraction among all four stages was tested by Chi-squared test for given probabilities, where control-edited melanosome counts have been considered as expected distribution. *P* values from Chi-squared test are shown in figure. Statistical significance of stage-wise comparison between gene knockouts and control-edited cells was done using one-sided 2-sample test of proportions.

To further confirm the validity of the screen results and gain insights into how select hits may affect melanosome biogenesis, we performed TEM imaging on the *WDR81, KLF6*, and *COMMD3* knockout (KO) cells and quantified melanosomes at various stages of maturation in comparison to control-edited and *TYR* KO cells (**Fig. 3C, D** and **S4C**). We found that *WDR81* KO cells had a significantly higher proportion of stage I melanosomes as compared to controls (one-sided 2-sample test of proportions, *P* = 1.41×10^−16^), as evidenced by presence of multiple vacuoles/vesicles (**Fig. 3D** and **Fig. S4C**). As stage I melanosomes are derived from vacuolar domains of the early endosomal compartments (*11, 19*), *WDR81* KO likely affects the early endosomal compartments. In agreement, a recent study showed that loss of *WDR81* caused an increase in endosomal PtdIns3P, leading to enlargement of early endosomes (*20*). Loss of *KLF6* displayed a significantly reduced proportion of stage IV melanosomes (one-sided 2-sample test of proportions, *P* = 9.15×10^−9^) (**Fig. 3D** and **Fig. S4C**), explaining the reduced pigmentation of the KO cells (**Fig. 3 A, B**) and suggesting that this transcription factor regulates expression of genes required for the final stage of melanosome maturation. Finally, deletion of *COMMD3* led to a significant increase in stage I and II non-melanized melanosomes (one-sided 2-sample test of proportions, *P* = 5.86×10^−8^), where fibrillar structure was visible, but transition to stage III/IV appeared to be defective (**Fig. 3D** and **Fig. S4C**). COMMD3 belongs to the evolutionary conserved COMMD (COpper Metabolism MURR1 or COMM domain) family of proteins, comprised of ten members. COMMD proteins homo/heterodimerize and associate with CCDC22 and CCDC93 proteins to form the CCC complex, involved in the recycling of copper-transporting ATPase 1 (ATP7A) (*18*). Both *CCDC22* and *CCDC93* as well as multiple *COMMD* genes (*COMMD2, COMMD3, COMMD4, COMMD5, COMMD8* and *COMMD10*) of the CCC complex were identified as hits in our screen. The key melanogenic enzyme, TYR requires copper as an essential cofactor and the lack of melanosome specific copper transporter ATP7A has been shown to result in severe pigmentation defects (*21*). Given that *COMMD3* KO phenotype is very reminiscent of *TYR* KO phenotype (**Fig. S1G** and **Fig. S4C**), it is tempting to speculate that COMMD3 and other CCC complex components may be involved in recycling of ATP7A copper transporter modulating TYR activity. Taken together, our screen identified novel genes involved in the regulation of melanosome biogenesis and maturation at different stages and through diverse biological mechanisms.

Next, we sought to understand the relevance of our CRISPR screen results for variation in human pigmentation. We first applied stratified LD score regression (S-LDSC) (*22*) to assess skin color heritability enrichment in the vicinity (within 100kb) of our screen hits, using publicly available summary statistics from a GWAS of European ancestry individuals in the UK Biobank. The 169 screen hits (at FDR <10%) were significantly enriched for skin color heritability (Fold-enrichment 9.61, standard error 3.80, *P* = 3.11×10^−2^) in comparison to all non-hits (FDR > 10%) (fold-enrichment 1.15, standard error 0.073, *P* = 3.37×10^−2^) (**Fig. 4A**). Next, we collected lists of genes implicated by various GWAS, including: a) all genes annotated to pigmentation or skin color-associated SNPs in the NCBI-EBI GWAS catalog, representing a mix of ancestries (*23*), b) genes nearby lead SNPs associated with skin color variation in white individuals from the UK Biobank, and c) genes nearby lead SNPs associated with pigmentation variation in African individuals (*24*). Relative to all other genes assayed, screen hits were strongly enriched for pigmentation genes across the three datasets (**Table S3**). For example, eleven of 169 hits lie within 100kb of a genome-wide significant lead SNP associated with skin color in UK Biobank individuals, compared to 266/20,119 non-hits (two-sided Fisher’s odds ratio 5.6, *P* = 2.75×10^−5^). Similar enrichments were observed for genes in the GWAS catalog (two-sided Fisher’s odds ratio 11.3, *P* = 6.13 x10^−6^) or nearby top pigmentation-associated SNPs in Africans (two-sided Fisher’s odds ratio 4.76, *P* = 2.1 x10^−4^). While some of the screen hits associated with GWAS peaks have known roles in pigmentation, others are novel. For example, of the screen hits in vicinity of the lead GWAS signals in the UK Biobank, three (*AP1G1, SLC12A9*, and *SLAIN1*) represent novel associations with melanogenesis (**Fig. S5**). These results indicate that both known and novel melanogenesis genes uncovered in our study contribute to skin color variation in human populations.

**Fig. 4.**
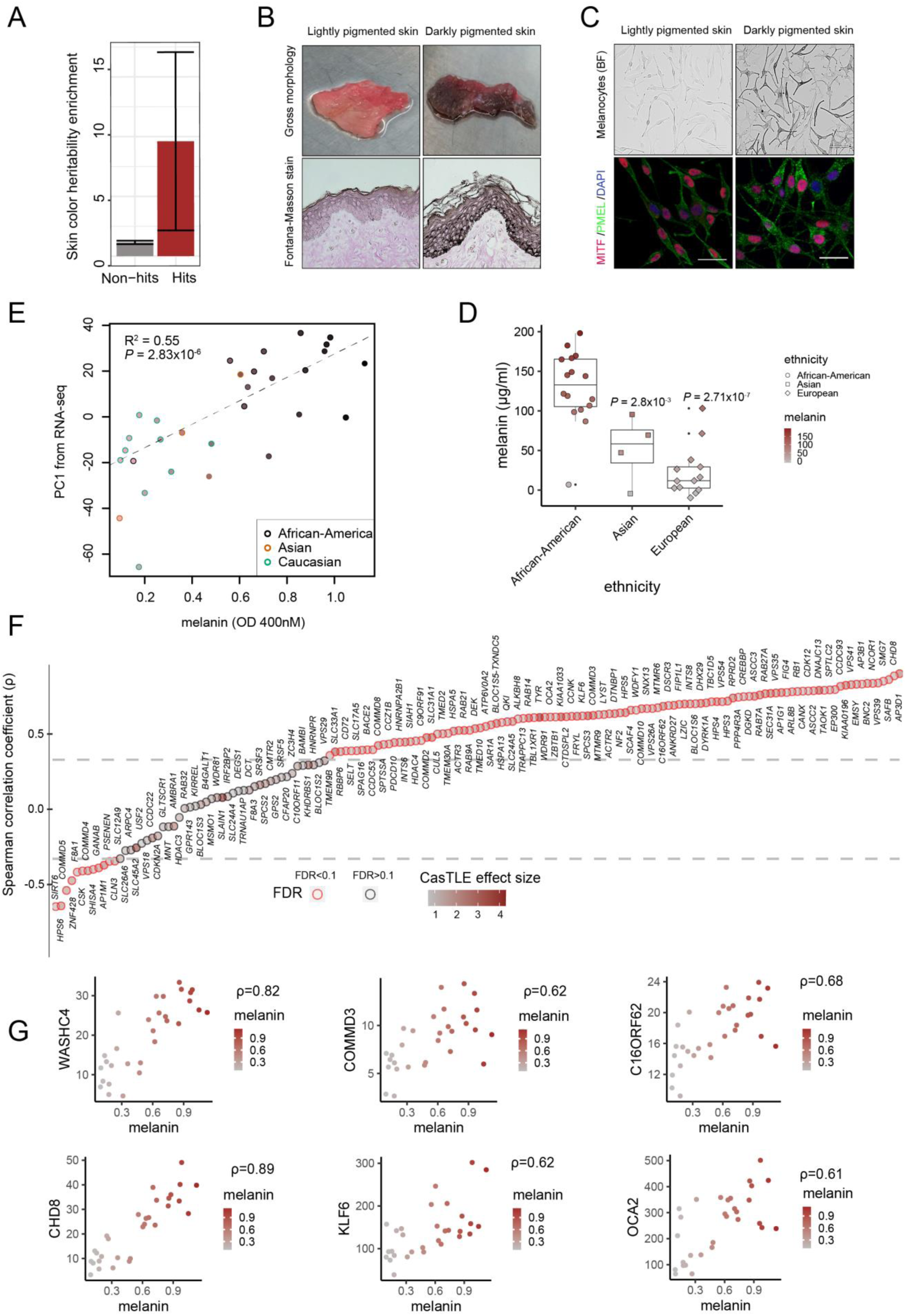
Pigmentation screen hits are differentially expressed in melanocytes of distinctly pigmented individuals and enriched for skin color heritability. **(A)** The effect size of skin color heritability enrichment (from UKBB GWAS) for the screen hits and non-hits is shown. The screen hits were significantly enriched for skin color heritability (Fold-enrichment 9.61, standard error 3.80, *P* = 3.11×10^−2^) in comparison to all non-hits (fold-enrichment 1.15, standard error 0.073, *P* = 3.37×10^−2^). Non-hits are defined as all genes assayed in the CRISPR screen with FDR > 0.1. The Error bars indicate 95% confidence interval for heritability fold-enrichment. (**B)** Gross morphology of foreskin tissue obtained from darkly and lightly pigmented human donors. Fontana-Masson staining shows melanin pigment (black color) in epidermis. (**C**) Brightfield images and immunofluorescence of MC obtained from darkly and lightly pigmented human donors confirms differential pigment levels and presence of MC markers MITF (red) and PMEL (green); DNA is stained with DAPI (blue). (**D)** Melanin quantification of MC derived from diversely pigmented individuals from different parent-reported ethnicities. Melanin was quantified spectrophotometrically by measuring OD (at 400nM wavelength) of cell lysates (2.5×10^5^ cells in 250 µl 1M NaOH with 10% DMSO) and comparing against a standard curve generated using synthetic melanin. Box plots show median log absorbance and IQR, whiskers are 1.5 × IQR. Significance tested by ANOVA followed by two-sided Welch t-test with Benjamini & Hochberg correction. *P* values shown are relative to African-American donors. (**E)** Correlation between RNA-seq gene expression profiles and melanin content of MC obtained from darkly and lightly pigmented human donors. Plotted is melanin concentration (OD 400nm) isolated from melanocytes from various donors versus first principal component of the RNA-seq analysis from the same melanocytes. Parent-reported ethnicity is indicated by the color of the outline of the plotting symbol. Fill color within each point represents measured melanin content. (**F)** Spearman’s rank correlation coefficient (*ρ*) comparing relationship between gene expression (TPM) of candidate genes identified as screen hits with melanin measurements in 30 MC samples of diverse skin color. Each point filled with brown color, intensity of which indicating CasTLE effect size. Horizontal lines indicate (−0.33, 0.33) correlation coefficient cut offs at 10% FDR. (**G)** Spearman’s correlation coefficient for selected CRISPR screen hits, with each point representing measurements from an individual donor. Plotted is melanin content (ordinate) vs RNA-seq expression level (TPM, abscissa). Scale bars (C, D): FM, BF, IF = 25 µM.

Our analyses thus far indicated that at least some melanin-promoting screen hits may be involved in regulating skin color variation in human populations. One of the key mechanisms by which phenotypic variation arises is through gene regulatory divergence (*25, 26*). We therefore reasoned that many pigmentation genes involved in skin color variation in humans are likely to be differentially expressed between lightly and darkly pigmented individuals. To survey transcriptomes of diversely pigmented human MC, we obtained foreskin tissues from 33 newborn males of diverse skin color and isolated primary MC from them (**Fig. 4B and Table S4**). Foreskin tissues were histologically analyzed for melanin presence in situ by Fontana-Masson (FM) staining, which confirmed the presence of differential melanin content in the epidermis of diversely pigmented individuals (**Fig. 4B**). Particularly, the basal epidermis of lightly pigmented skin displayed fainter FM staining than darkly pigmented skin. Isolated MC all expressed key lineage transcription factor MITF and melanosome specific marker PMEL, but had distinct pigmentation levels (**Fig. 4C**). We spectrophotometrically quantified total melanin content of isolated MC by measuring absorbance of MC lysates at 400nM and comparing them against a standard curve generated using a synthetic melanin (**Fig. 4D and Fig. S1D**). As expected, melanin content correlated with the parent-reported donor ancestry, with MC derived from African-American donors showing higher melanin content, followed by Asian and European donors, respectively (**Fig. 4D, Table S4**).

Next, we performed RNA-seq on isolated diversely pigmented MC samples (n=30) (**Table S5**). Principal component analysis (PCA) of RNA-seq results demonstrated that gene expression divergence correlated well with measured melanin content of the cells, even more so than reported ethnicity (**Fig. 4E**). We then calculated Spearman’s rank correlation coefficients (ρ) between expression (TPM) of genes corresponding to our screen hits and melanin content (OD at 400nM). Out of the total 169 screen hits, 158 genes had measurable expression in all MC, and were therefore used for these calculations. We found that the expression of 107 hits (out of 158) was significantly positively correlated (q value < 0.1) with melanin, 12 genes were negatively correlated (q value < 0.1) and 39 were not significant (q value > 0.1) (**Fig. 4F, G, Fig. S6 and Table S6**), demonstrating that the majority of screen hits are more highly expressed in darkly pigmented MC. The key known skin variation genes picked up by our screen, such as *OCA2* and *SLC24A5*, as well as genes encoding BLOC and AP complexes involved in melanogenesis, showed significant upregulation in dark MC (with a notable exception of BLOC-2 subunit HPS6, which showed negative correlation) (**Fig. 4F, G, Fig. S6** and **Table S6**). Importantly, endosomal recycling complexes identified here, including components of retromer, retriever, WASH, ARP2/3, and CCC complexes were also differentially expressed between lightly and darkly pigmented MC (**Fig. S6** and **S7**). At the gene regulatory level, chromatin remodelers and modifiers (*EP300, CHD8*, and *CREBBP*), transcriptional repressor complex (*NCOR1, HDAC4* and *TBL1XR1*), alternative mRNA splicing regulators (*QKI, HNRNPA2B1*, and *DYRK1A*) and RNA 3’-end processing factors (*INTS6, INTS8* and *FIP1L1*) were all significantly upregulated in dark MC (**Fig. 4G, S6** and **S7**). Together, these results indicate that the majority of novel melanogenic genes identified in our screen are differentially expressed among distinctly pigmented individuals, and thus may play a role in mediating human skin color divergence.

In this study, we demonstrated that changes in melanosomal composition can be robustly quantified by measuring the side scattering property of pigment cells. By exploiting this relationship, we directly surveyed genes involved in melanosomal biogenesis/maturation without being confounded by the indirect factors influencing pigmentation through impacting melanocyte development, survival, migration and other processes. Our CRISPR screen uncovered a wealth of new regulators of melanogenesis involved in diverse molecular functions, such as transcriptional and posttranscriptional gene regulation, vesicle organization and intracellular transport, the latter including several molecular complexes involved in endosomal recycling. While the key role of early endosomes in melanogenesis is well established, our work implicates new cargo recycling pathways in melanosome function. These candidates and pathways merit further exploration in the context of disease, as aberrant melanin production is associated not only with skin pigmentation disorders, but also with Parkinson’s disease and auditory disorders, which have been linked to loss of neuromelanin in *substantia nigra* and cochlear melanin in the inner ear, respectively (*27, 28*).

Importantly, the majority of novel melanogenesis genes uncovered in our study are more highly expressed in MC of darkly pigmented individuals, which is in agreement with their melanin-promoting function and suggests that regulatory divergence in these genes contributes to the normal range skin color variation in humans. At least some of the transcriptional differences linked to the melanin content must be genetically encoded (as opposed to being a consequence of distinct cellular environments of lightly and darkly pigmented MC), because we see significant skin color heritability enrichment in the vicinity of our screen hits. Therefore, in addition to the aforementioned insights into melanosome biology and disease, results presented here will provide a rich resource for further studies of genetic architecture and evolutionary history of skin color variation in humans.

## Materials and Methods

### Cell culture

All cells were routinely tested for mycoplasma contamination using MycoAlert detection kit (Lonza).

#### Pluripotent stem cells (PSC) culture

The SOX10::GFP reporter H9 human embryonic stem cells (hESC) were initially cultured on mouse embryonic feeder cells as described previously (*29*) and gradually adapted to feeder-free culture. H9 hESC SOX10::GFP reporter line and H20961 human induced pluripotent stem cells (iPSCs) were cultured in feeder-free, serum-free medium mTESR-1 (StemCell technologies).

Both pluripotent stem cells (PSC) i.e. H9-hESC SOX10::GFP reporter and H20961 iPSC were passaged ∼1:6 every 4-5 days by incubating them in ReLeSR (StemCell technologies) for 1 min at room temperature. The ReLeSR was then aspirated and the culture plates were incubated for 6-7 mins at 37°C. mTESR-1 was added to the cells and the plates were gently tapped to detach the cells, which were triturated and re-plated on tissue culture dishes coated overnight with growth-factor-reduced Matrigel (BD Biosciences).

#### PSC differentiation into melanocytes

The SOX10::GFP reporter H9 hESC and H20961 iPSCs were differentiated into melanocytes following a previously published protocol with some modifications (*7*). Cells were plated on Matrigel coated plates in mTESR-1 medium and when colonies reached 70% confluency, the medium was changed to differentiation medium composed of KSR medium (1x knockout DMEM, 15% knockout serum replacer, 1xNEAA, 1x Glutamax, 55 mM 2-mercaptoethanol (Gibco) containing 10 μM SB431542 (Selleck Chemicals) and 500nM LDN193189 (Selleck Chemicals) for 48 hours with medium change every 24 hours. After 48 hours, in the above medium, 3 μM CHIR99021(Selleck Chemicals) was added. After 72 hours, cells were fed with KSR media containing 10 μM SB431542 and 3 μM CHIR99021. On day 4 (after 96 hours), cells were fed with 75% KSR, 25% N2 medium plus 3 μM CHIR99021. The N2 medium is comprised of: 1x DMEM-F12 (Life Technologies),1x N2 NeuroPlex supplement (GeminiBio), 5.76 × 10^−5^ mg/ml progesterone (Sigma), 1.55 mg/ml D-glucose (Sigma), 1x 55 mM 2-mercaptoethanol (Gibco), 1x antibiotic-antimycotic (Gibco)). On day 6, cells were fed with 50% KSR, 50% N2 media containing 3 μM CHIR99021 plus 25 ng/ml BMP4 and 100 nM endothelin-3 (Alfa Aesar). On day 8, media was switched to 25% KSR media, 75% N2 media plus 3 μM CHIR99021, 25 ng/ml BMP4 (PeproTech), and 100 nM EDN3. On day 10, cells were fed with 100% N2 media containing 3 μM CHIR99021, 25 ng/ml BMP4, and 100 nM EDN3. On day 11, cells were treated with Accutase for 30 mins and washed twice with neurobasal medium (Life Technologies), counted, resuspended at a concentration of 2×10^6^ cells/ml and plated on poly-L-ornithine, laminin, and fibronectin coated dishes as 20 μl volume droplets. Droplets were incubated at 37°C for 20 mins followed by addition of MEL medium (50% neurobasal, 30% low glucose DMEM (Life Technologies), 20% MCDB201 (Sigma), 1x antibiotic-antimycotic (Gibco), 1x Glutamax (Gibco), 1x ITS supplement (BD), 100uM ascorbic acid (Sigma), 50ng/ml cholera toxin (Sigma), 50ng/ml StemCellFactor (Fisher Scientific), 50nM dexamethasone (Sigma), 100nM EDN3, 2% Gem21 Neuroplex (geminiBio), 3 μM CHIR99021, 4ng/ml FGF2 (PeproTech), 500um dbcAMP (Sigma), 25ng/ml BMP4). MEL medium was changed every 3 days and cells were passed every 5-6 days. Cells started showing pigmentation past day 25 of differentiation. As the cells differentiated from MB to MC state, cells were analyzed for side scatter (SSC) changes by FACS as well as melanin quantification by measuring OD at 400nM and comparing against a standard curve generated using synthetic melanin.

#### Primary human melanocytes culture

Foreskin tissues were procured from 1-2 day old newborn males of diverse ethnicity following Institutional IRB protocol guidelines. Skin tissues were processed as described previously (*30*). Briefly, tissues were washed 3 times with 1x PBS, cut into pieces (∼5mm×5mm). A piece was fixed in 10% neutral buffered formalin and provided to Stanford Pathology core for Fontana-Masson staining. The rest of the tissue was enzymatically digested with dispase I (Zen-Bio) for 15-20 hours at 4°C. Afterwards, the epidermis was peeled from the dermis manually using fine forceps. The epidermis was treated with trypsin-EDTA (Life Technologies) for 15-20 mins at 37°C with intermittent trituration. The cell suspension was neutralized using soy trypsin neutralizing solution (Life Technologies), filtered through a 70 μm cell strainer (BD Biosciences), centrifuged at 200g for 5 min, and cultured in the presence of serum-free medium 254 supplemented with 1x human melanocyte growth supplement (HMGS) for 72 hours without medium change. Afterwards, fresh medium was replenished every other day.

#### Melanoma cell line culture

MNT1 cells were cultured in high glucose DMEM (Life Technologies) supplemented with 10% fetal bovine serum, 1x antibiotic-antimycotic and 1x Glutamax.

### Plasmids, SpCas9 expression, gene knockout and rescue cell lines generation

Pb-TRE3G-SpCas9-IRES-Blast, a piggyBac doxycycline inducible SpCas9 expression vector (see image below), initially generated in Wysocka laboratory was used in this study.

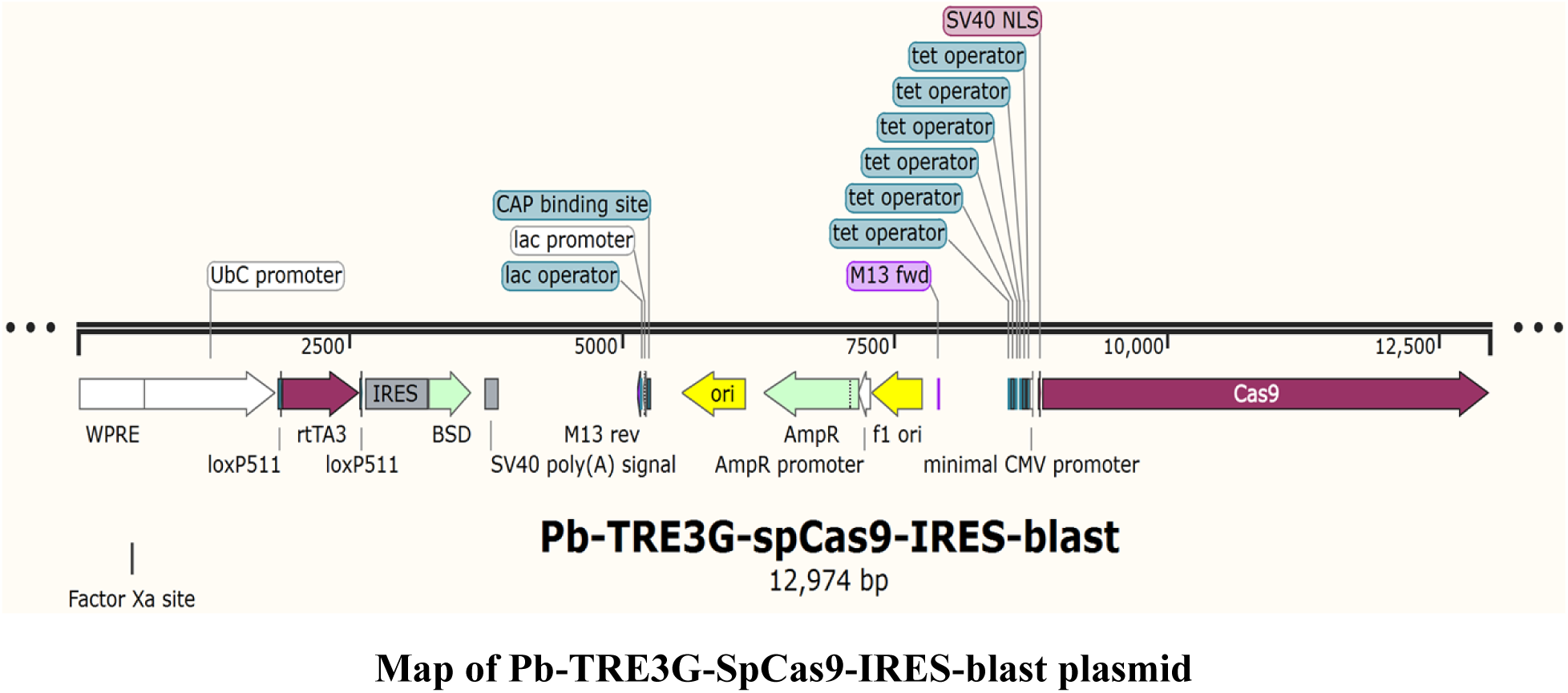

This plasmid efficiently expressed SpCas9 under a doxycycline inducible promoter and also contained a blasticidin selection cassette and was subsequently used for creating the Cas9-MNT1 cell line that was used for both genome-wide screening as well as screen hit validation experiments. To generate Cas9-MNT1 cell line, MNT1 cells were transfected with Pb-TRE3G-SpCas9-IRES-Blast and piggyBac transposase (PB210PA-1, System Biosciences) in 1:3 ratio using lipofectamine (Life Technologies) following the manufacturer’s instructions. Cells were selected with 5µg/ml blasticidin treatment for a week. Afterwards, single cells were plated to generate a clonal cell line and the efficient induction of SpCas9 nuclease from the clonal cell line was confirmed by western blot.

In the screen hit validation experiment, the Pb459-U6-sgRNA-EF1a-PuroR vector used for sgRNA expression in the Cas9-MNT1 cells was designed in two steps. First, the lentiviral backbone of the pMCB320 vector was replaced with the piggyBac backbone from the PB-TRE-dCas9-VPR (Addgene, 63800) vector using EcoRI and XbaI restriction enzymes. Next, PuroR-T2A-mCherry was replaced with the PuroR cassette using NheI and EcoRI. Pairs of oligonucleotides carrying sgRNA sequences (see table below) and the BlpI and BstXI overhangs were phosphorylated and annealed. The annealed oligos were subsequently ligated into the BlpI and BstXI digested, and gel purified, Pb459-U6-sgRNA-EF1a-PuroR backbone.

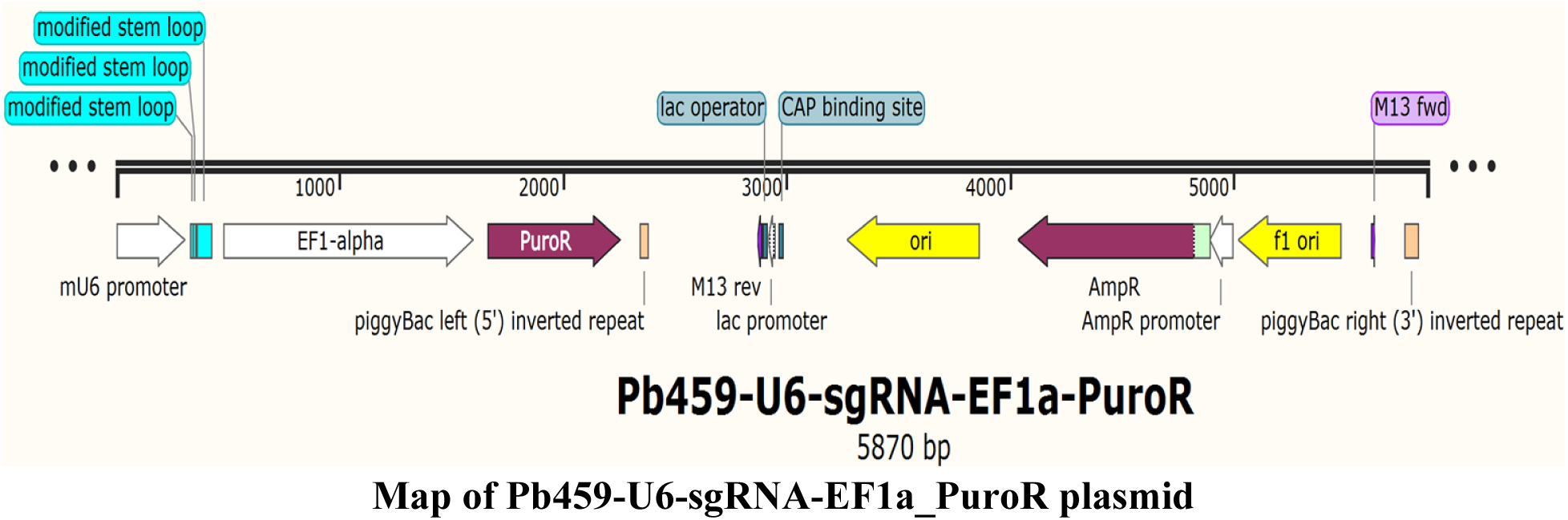

**Table.**
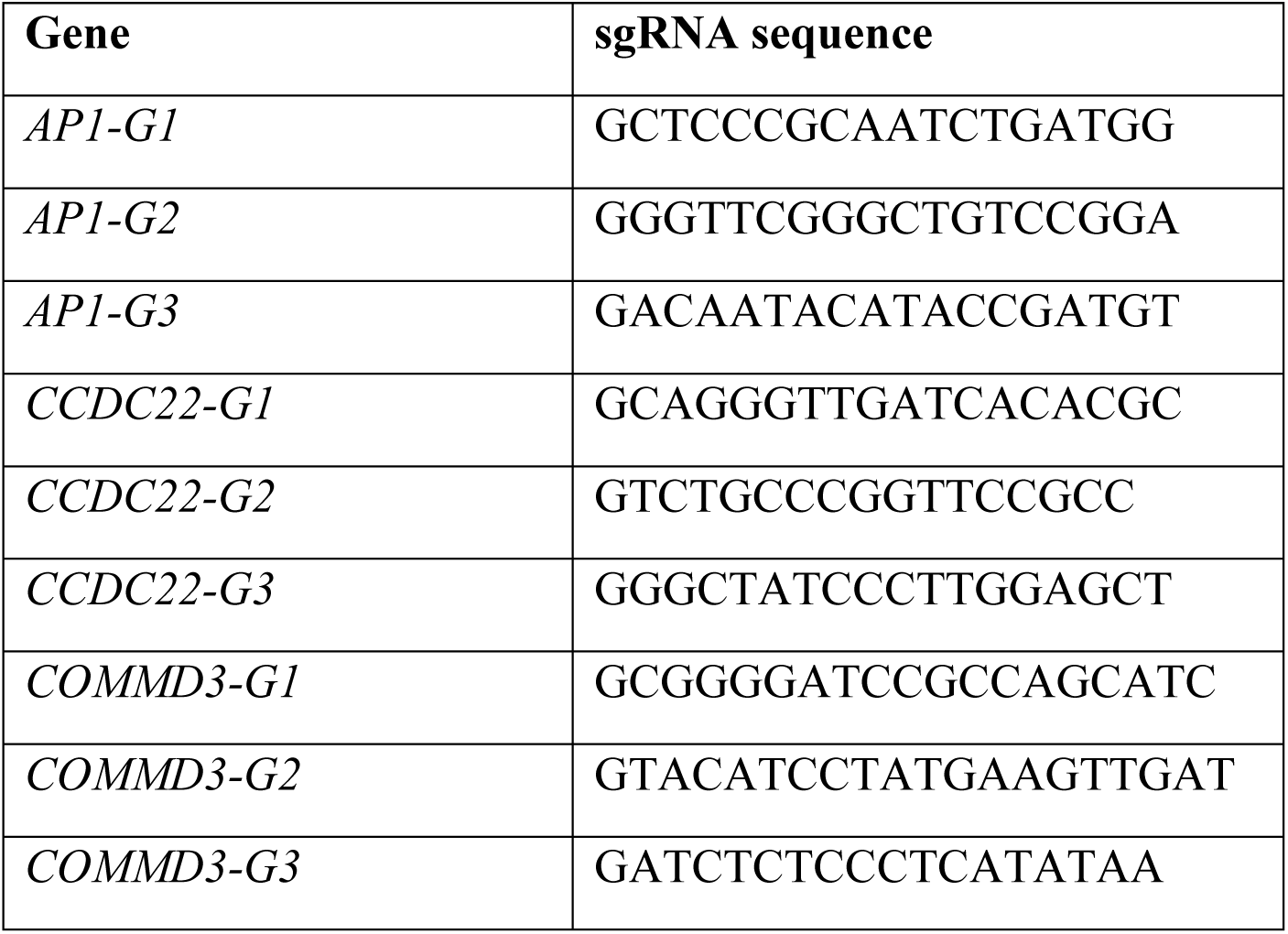
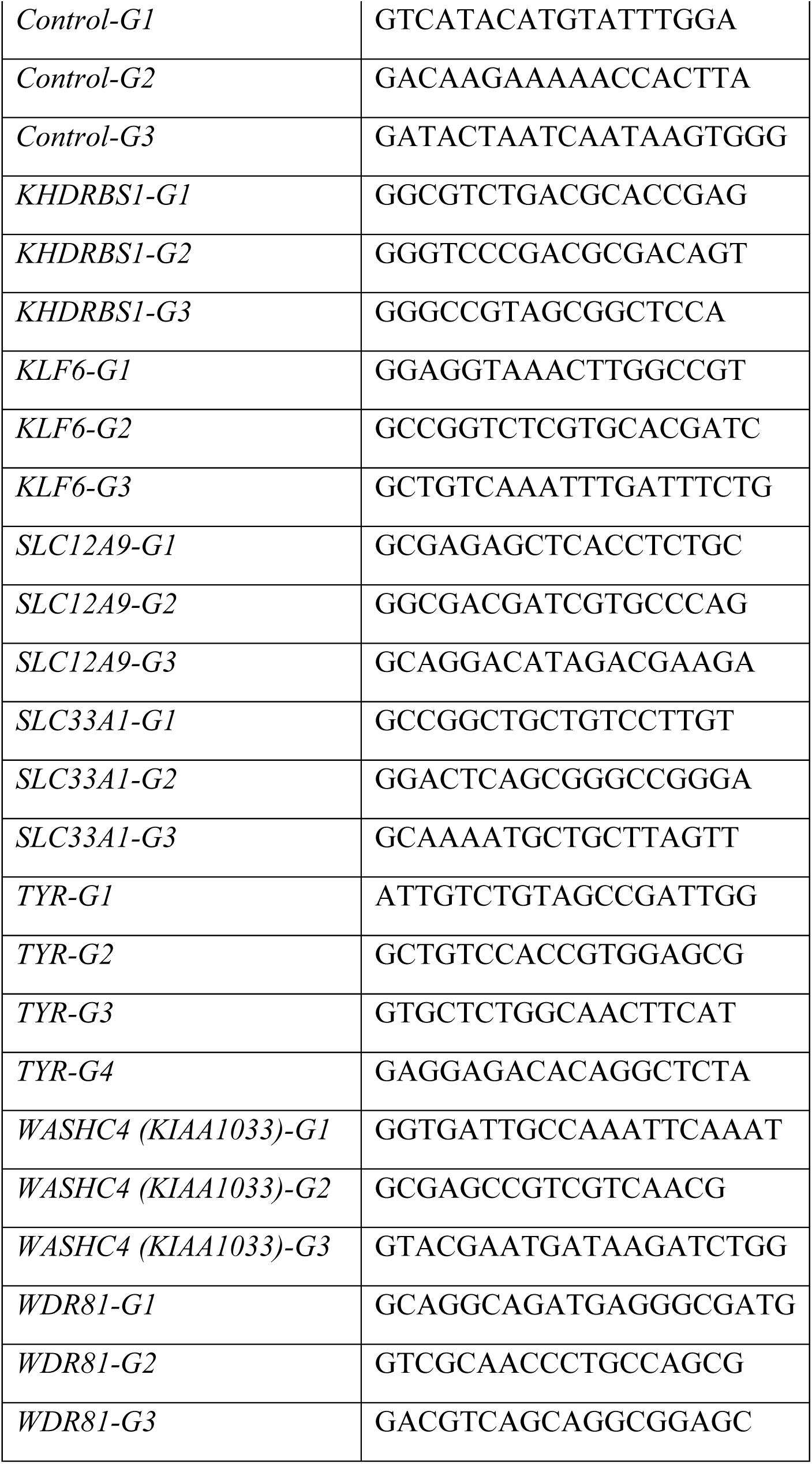
sgRNA sequence used for knockout experiments.

After cloning the Pb459-U6-sgRNA-EF1a-PuroR vector expressing unique sgRNAs, Cas9-MNT1 cells were transfected with Pb459-U6-sgRNA-EF1a-PuroR and piggyBac transposase (PB210PA-1, System Biosciences) in a 1:3 ratio using lipofectamine (Life Technologies) following manufacturer’s instructions. Transfected cells were selected with 1µg/ml puromycin treatment for 4-6 days. Afterwards, cells were treated with 2µg/ml doxycycline for 2 weeks followed by cell lysis, melanin quantification and FACS analysis on FACSAria (BD)/ CytoFLEX (BD) for SSC measurements. FACS analyses were done using FlowJo software (BD) and flowCore() package in R.

To generate the *TYR* expression plasmid used in the tyrosinase rescue studies, a wild copy of the *TYR* gene was PCR amplified form the pEGFP-TYR plasmid (Addgene, 32781) with NheI and AgeI overhangs. The amplicon was then digested and ligated to the already digested and gel purified PB-TRE-dCas9-VPR plasmid (Addgene, 63800) backbone. The resulting vector Pb-TRE3G-TYR-HygR (see image below) contained the TYR cassette under the doxycycline inducible TER3G promoter and also contained a hygromycin selection cassette.

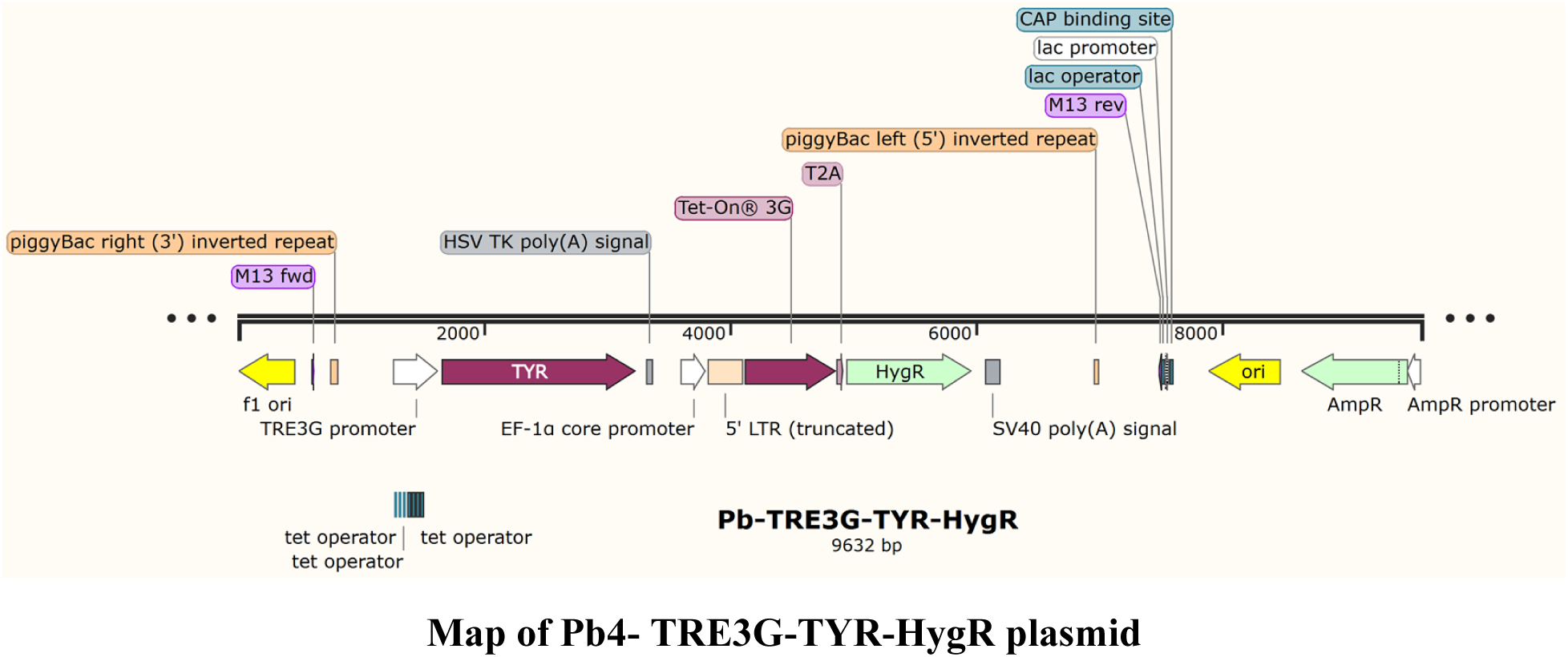

*TYR-G1-G4* knockout cell lines were transfected with the Pb-TRE3G-TYR-HygR plasmid and piggyBac transposase (PB210PA-1, System Biosciences) in a 1:3 ratio using lipofectamine (Life Technologies) following the manufacturer’s instructions. Cells were selected with 40µg/ml hygromycin for a week followed by 2µg/ml doxycycline treatment for a month to re-express *TYR* on *TYR* knockout cell lines. Afterwards, cells were lysed for melanin quantification and SSC measurements.

### Total cellular melanin quantification

Total melanin quantification was done as described previously (*31*). For primary human melanocytes and PSC-derived melanocytes, 2.5×10^5^ cells were counted, centrifuged at 4000xg for 5 min and washed twice with PBS. Then the cell pellet was dissolved in 250 μl 1M NaOH with 10% dimethyl sulfoxide (DMSO). For the MNT1 melanoma cell line, 6.0×10^5^ cells were dissolved in 120μl 1M NaOH with 10% dimethyl sulfoxide (DMSO). Cell lysate was incubated at 80°C for 2 hours. Afterwards, 100 μl of lysate was placed in a 96 well plate to measure absorbance at 400nM wavelength on an Infinite 200 Pro plate reader (Tecan). A standard curve was generated using synthetic melanin (Sigma) and melanin concentration was determined by reading OD 400nM against the standard curve.

### Immunocytochemistry

Immunocytochemistry was performed as described previously (*30*). Briefly, cells were fixed in 4% paraformaldehyde, permeabilized with PBS with 0.1% Triton X-100 and blocked with 1%BSA/0.01% Triton X-100. Afterwards, cells were incubated with one of the following anti-human primary antibodies diluted in blocking buffer: goat PAX3 (1:100; 4°C overnight; Santa Cruz, sc-34916), mouse TYRP1 (MEL-5) (1:500; 4°C overnight; BioLegend, 917801), rabbit MITF (1:200; 4°C overnight; Abcam,ab59232), goat SOX10 (1:50; 4°C overnight; Santa Cruz, sc-17342), mouse HMB45 (1:20; 4°C overnight; Life Technologies, 081050). Afterwards, cells were incubated with anti-mouse or anti-goat Alexa Fluor 488 and Alexa Fluor 594 antibodies (1:400; 1 hr at RT; Invitrogen) and counter-stained with DAPI nuclear dye (0.5 µg/ml in PBS; 10 min; Sigma). Cells incubated with secondary antibodies alone served as negative controls. Images were acquired on either an inverted microscope (Nikon Ti2) or on a confocal microscope (Leica TSC SP2).

### Western blotting

Cas9-MNT1 cells were induced for 48 hours with 2µg/ml doxycycline to express the Cas9 nuclease. Afterwards, whole cell extracts were obtained by lysing cells in ice cold lysis buffer (300 mM NaCl, 100 mM tris (pH 8), 0.2 mM EDTA, 0.1% NP-40, 10% glycerol, 1 mM phenylmethylsulfonyl fluoride (PMSF), and protease inhibitor cocktail (complete mini, Roche Applied Sciences, IN)) for 20 mins at 4°C. The supernatant was collected, the protein concentrations determined by Bradford assay using Bradfrod reagent (Bio-Rad) and serial dilutions of proteins were separated on an SDS–polyacrylamide gel electrophoresis (SDS-PAGE) gel. Afterwards, proteins were transferred to nitrocellulose membranes and immunoblotted using mouse monoclonal Cas9 antibody (7A9-3A3, Cell Signaling Technology), HRP-conjugated rabbit monoclonal GAPDH antibody (D16H11, Cell Signaling Technology) and HRP-conjugated mouse monoclonal anti-beta actin antibody (AC-15, Abcam). Protein bands were detected using Lumi-Light (Roche) chemiluminescence kit as per the manufacturer’s instructions.

### Transmission election microscopy

Cells were culture on ACLAR coverslips and fixed with 2% glutaraldehyde and 4% paraformaldehyde in 0.1M Sodium Cacodylate buffer, pH 7.4, for 1 hour at room temperature. Thin sections were cut and post-fixed with 1% Uranyl acetate. Sections were subsequently dehydrated and Epon infiltrated, and examined with JEOL JEM-1400 series Transmission Electron Microscope (TEM) at Stanford Cell Science Imaging Facility. TEM images were acquired for different gene knockouts as well as control-edited cells and melanosomes at various stages of maturation, i.e. stage I-IV were counted in a blinded experiment where the person doing the counting was unaware of the experiment design and expected outcome.

### Genome-wide CRISPR screen

MNT1 cells were transfected using Lipofecatime (Life Technologies) with piggyBac transposase (PB210PA-1, System Biosciences) and a piggyBac plasmid expressing SpCas9 under a doxycycline inducible promoter and containing a blasticidin selection cassette. Clonal cells were selected with 5µg/ml blasticidin treatment for a week. These cells, termed as Cas-MNT1 cells, expressed SpCas9 upon doxycycline treatment and were used for the genome-wide CRISPR screen.

The pMCB320 vector based 10-sgRNA per gene CRISPR-Cas9 library base has been described previously (*14*). This library consists of nine sub-libraries containing ∼ 200,000 guides targeting over 20,000 genes and 13,500 control guides. Control guides included non-targeting guides (i.e. no binding sites in the genome) and safe-targeting control guides targeting genomic locations with no annotated functions (*14*).

A large batch of nine CRISPR-Cas9 lentiviral sub-libraries was packaged using methods described previously with some modifications (*29*). Briefly, 293T packaging cells were transfected with CRISPR-Cas9 sub-libraries plasmid and a third-generation lentiviral packaging mix (1:1:1 mix of pMD2.G, pRSV-Rev, and pMDLg/pRRE) in a 1:1 ratio using polyethylenimine. After 12 hours of transfection, fresh medium (high glucose DMEM plus 10% fetal bovine serum) was changed to the 293T cells and subsequently two batches of lentivirus were collected at 24 hours and 48 hours. These two batches were combined, passed through 0.45 um syringe filter (Millipore) and ultracentrifuged (50,000xg at 4C for 2 hours) and viral pellet was resuspended in opti-MEM medium and stored in −80C until further use. Cas9-MNT1 cells were transduced with serially diluted lentiviral sub-libraries and FACS analyzed for mCherry positive cells to determine transduction efficiency and multiplicity of infection (MOI) calculations. For two independent genome-wide screens, an MOI of 0.1 was used to infect ∼200×10^6^ Cas9-MNT1 cells. Cells were selected with 1µg/ml puromycin for two weeks to achieve more than 90% mCherry positive cells and then were treated with 2µg/ml doxycycline for two weeks for Cas9 induction and subsequent gene editing. At this point, cells from the top and bottom 10% of the SSC distribution were FACS sorted on FACSAria cell sorters (BD). For each replicate, ∼50×10^6^ low SSC and ∼50×10^6^ high SSC fractions were sorted, totaling ∼100×10^6^ cells at 500x coverage. Genomic DNA was isolated from 200×10^6^ cells using QIAamp DNA blood midi kit (Qiagen) followed by DNA purification using DNeasy PowerClean Pro Cleanup kit (Qiagen) to remove coprecipitated melanin following the manufacturer’s instructions.

PCR amplification of inserted sgRNAs from the genomic DNA was done with the Herculase II Fusion DNA Polymerase (Agilent Technologies) using the primers oMCB-1562 and oMCB-1563. All of the genomic DNA was PCR amplified using the following PCR protocol: 1× 98°C for 2 min, 18× 98°C for 30 sec, 59.1°C for 30 sec, 72°C for 45 sec, 1× 72°C for 3 min. Each PCR reaction included 300 ng of genomic DNA, 20 μL 5X Herculase buffer, 2.5 μL of 10 mM dNTPs, 1 μL 100 μM oMCB-1562, 1 μL 100 μM oMCB-1563, 0.5 μL Herculase II Fusion DNA polymerase, and water to make up the volume to 100 μL. PCR amplicons from each PCR reaction for each group were pooled together and a second five step PCR reaction was set up for each group as follows: 5 μL amplicon from PCR reaction 1, 20 μL 5X Herculase buffer, 2.5 μL of 10 mM dNTPs, 1 μL 100 μM oMCB-1439, 1 μL 100 μM of a unique barcoded primer, 0.5 μL Herculase II Fusion DNA polymerase, and water to make up the volume to 100 μL. PCR conditions for the second PCR reaction were: 1× 98°C for 2 min, 19× 98°C for 30 sec, 59.1°C for 30 sec, 72°C for 45 sec, 1× 72°C for 3 min. PCR amplicons from each group were pooled, and 50 μL were run on a 1% TAE-agarose gel. PCR bands were excised, and purified using a Qiagen Gel Purification Kit as per manufacturer’s instructions. The sgRNA libraries were sequenced by deep sequencing on an Illumina Hi-Seq using the custom sequencing primer oMCB-1672. Computational analysis of the screen was done using the CasTLE maximum likelihood estimator, which utilizes the background of negative control guides RNAs as a null distribution model against which gene effect sizes, confidence scores and *P* values are determined. We used safe-targeting guides as control guides in our analysis. The results of the screen are shown in table S1. GO term enrichment analysis of CRISPR screen hits was performed using Gene ontology tools GOrilla (*32*) and REVIGO (*33*). Screen hits at 10% FDR were used as input gene set in GOrilla for GO term enrichment analysis for biological processes and components. The enriched GO terms with FDR corrected *P* value <0.05 were then used in REVIGO to remove the redundant GO terms and visualize as graphs (**Fig. S3**).

### RNA-seq

Total RNA was isolated from 1×10^6^ primary melanocytes (MC) using an RNA isolation kit (Zymo Research) as per the manufacturer’s instructions. mRNA was isolated using two rounds of Dynal oligo(dT) beads (Invitrogen) purification of 10ug total RNA. The mRNA was fragmented with 10X Fragmentation Buffer (Ambion) for 5 minutes and afterwards purified. First strand cDNA synthesis was carried out using Random Hexamer Primers (Invitrogen) and the SuperScript II enzyme (Invitrogen) followed by second strand cDNA synthesis with RNase H (Invitrogen) and DNA Pol I (Invitrogen). The cDNA was purified with QIAquick column (Qiagen). Sequencing libraries were prepared using Ovation Ultralow System (Nugen/Tecan) following manufacturer’s instructions. Sequencing adaptors were cleaned by magnetic beads (Agencourt XP) size selection. Libraries were QCed by Qubit and Bioanalyzer (Agilent) and multiplexed in six to eight samples per lane for single-end or paired-end sequencing on Illumina HiSeq 2500 or NEXTseq platforms.

Reads were aligned to hg38 genome model with hisat2 and an ab initio transcript discovery was performed using stringtie. These gene models were reconciled with human gencode 29 gene models using gffcompare. Read coverage over the models were obtained with featureCounts. Expression values as approximate TPMs were calculated at the transcript level without library fragment length correction and subsequently summed for each gene, if multiple transcripts per gene were present. Log TPM + 1 expression values for 2216 most variable genes were used to calculate PCA for exploratory analysis. For gene expression (TPM) and melanin (OD 400nM) correlation analysis, Spearman’s rank correlation coefficients were calculated using cor() function and *P* values were obtained by corr.test() from library(‘psych’) in R. Obtained *P* values were further FDR corrected with qvalue().

### Skin color heritability enrichment analysis

White British UK Biobank GWAS summary statistics were obtained from the Neale lab website (http://www.nealelab.is/uk-biobank/). GWAS was clumped into lead SNPs at various significance thresholds (5e-8, 1e-6, 1e-4) using plink v1.9, with the following parameters: -- clump-p1 (*6*) --clump-p2 (*6*) --clump-r2 0.01 --clump-kb 10000. A White British-specific LD reference panel from Sinnott-Armstrong and Naqvi, 2020 (*34*) was used. NCBI-EBI GWAS Catalog results were accessed in May 2020; all associations with the “skin pigmentation” search term were downloaded, and all genes in the “Mapped gene” column were used as the gene set. Since the top 1000 skin color SNPs in African populations were not pruned or clumped for independence, we primarily assigned each SNP to the closest gene. For stratified LD score regression analyses, annotations were constructed using 100kb up- and downstream of either CRISPR activators, repressors, or both. Coefficient Z-scores and heritability enrichments were calculated relative to the set of baselineLD annotations provided by Finucaine et al. (*22*).

## Acknowledgments

We sincerely thank Prof. Michael S. Marks (University of Pennsylvania) for providing MNT1 cells, Prof. Lorenz Studer (Memorial Sloan Kettering Cancer Center) for H9 SOX10::GFP reporter cell line and Prof. Yoav Gilad (University of Chicago) for H20961 iPSC. We thank John Perrino of Stanford Cell Science Imaging Facility for his help with transmission electron microscopy. We thank members of the Wysocka lab for comments on the manuscript; Deepika Verma for help with schematic figures; Funding: This work was supported in part by the Howard Hughes Medical Institute, NIH R35 GM131757 and Ludwig Center for Cancer Stem Cell Research and Medicine and Stinehart-Reed award (to J.W.), and Department of Defense grant CA160997 (to V.K.B.).

## Author contributions

V.K.B. conceived and designed the study, performed experiments, analyzed data including bioinformatics analyses of CRISPR screen, and wrote the manuscript with input from all co-authors; V.K.B., performed FACS experiments with help from M.A., N.A. and T.C.K.; T.C.K. did blinded melanosome counting of TEM images; J.T., M.B. provided advice and technical support with CRISPR screening; J.M. provided advice on data analyses; .S.N. performed heritability analyses of screen hits; J.K.P. provided advice on heritability analyses; T.S. performed bioinformatics analyses, provided advice on data analyses and experimental design; J.W. conceptualized and supervised the project and co-wrote the manuscript with V.K.B.

## Data and materials availability

All data needed to evaluate the conclusions in the paper are present in the paper, Supplementary Materials, and/or Genomic datasets available in the Gene Expression Omnibus (accession number pending).

## Supplementary Materials

**Supplementary figure legends S1-S7**

**Supplementary table legends S1-S6**

**Fig. S1.**
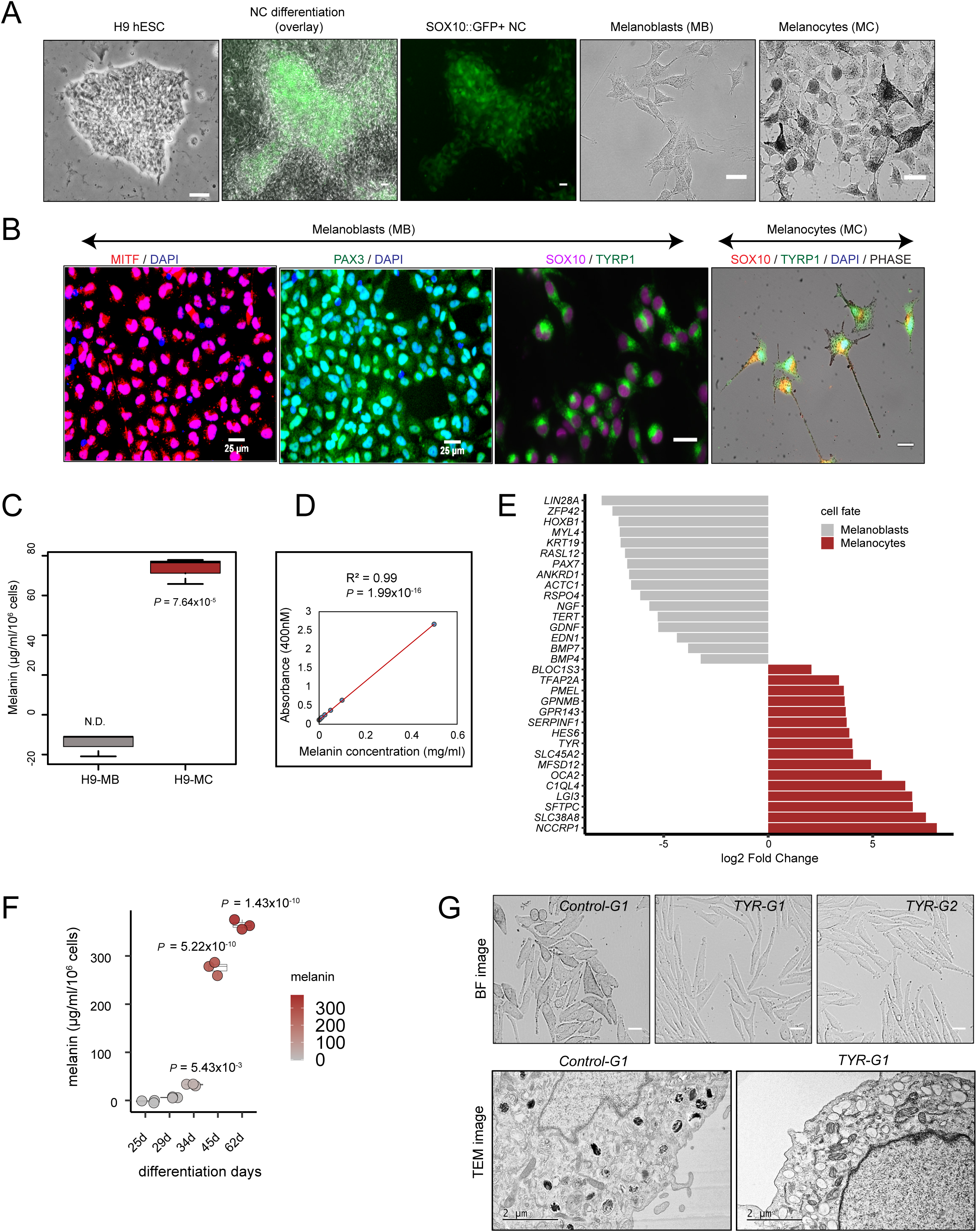
Derivation of melanoblasts and melanocytes from pluripotent stem cells and melanin concentration measurements. (**A)** Stepwise differentiation of H9 human embryonic stem cells (hESC) *SOX10::GFP* reporter line into neural crest (NC) as shown by GFP+ cells, non-pigmented melanoblasts (MB) and finally, pigmented melanocytes (MC). (**B)** Expression of select markers during differentiation is shown: for MB fate immunostainings for MITF, PAX3, SOX10 and TYRP1; for MC, immunostainings for SOX10 and TYRP1 overlapping with phase contrast image. (**C)** Melanin content of H9 hESC-derived MB and MC was determined spectrophotometrically by measuring OD (at 400nM wavelength) of cell lysates (2.5×10^5^ cells in 250 µl 1M NaOH with 10% DMSO) and comparing against a standard curve generated using synthetic melanin. Box plots show median and IQR, whiskers are 1.5 × IQR (n=3). Significance tested by Welch t-test. **(D)** Standard curve depicting relationship between melanin concentration and absorbance at 400nM. R^2^ and *P* value determined by linear regression. (**E)** In comparison to MB, MC express mature melanogenic genes. The first five genes with highest log2 fold changes are the 5 most differentially regulated genes in each category. Log2 fold changes calculated from RNA-seq analysis of four MB and three MC cell populations derived from H9 and H20961 pluripotent stem cells. (**F**) Melanin content at different days of differentiation from hESC to melanocytes was determined spectrophotometrically by measuring OD (at 400nM wavelength) of cell lysates (2.5×10^5^ cells in 250 µl 1M NaOH with 10% DMSO) (n=3 differentiations) and comparing against standard curve generated using synthetic melanin as shown in (D). Central value is median. Box plots show median and IQR, whiskers are 1.5 × IQR. Significance tested by ANOVA followed by two-sided pairwise Welch t-Test with Benjamini & Hochberg correction. *P* values shown are relative to day 29 of differentiation. (**G**). Brightfield and TEM images of *TYR* knockout in MNT1 cells. TEM images shows predominance of stage I-II melanosomes and lack of mature melanosomes upon TYR loss. (n = 4 sgRNAs treatments). Scale bars: A, B, G (BF images) = 25µM. TEM Scale bars: 2 µM. N.D. = not detected.

**Fig. S2.**
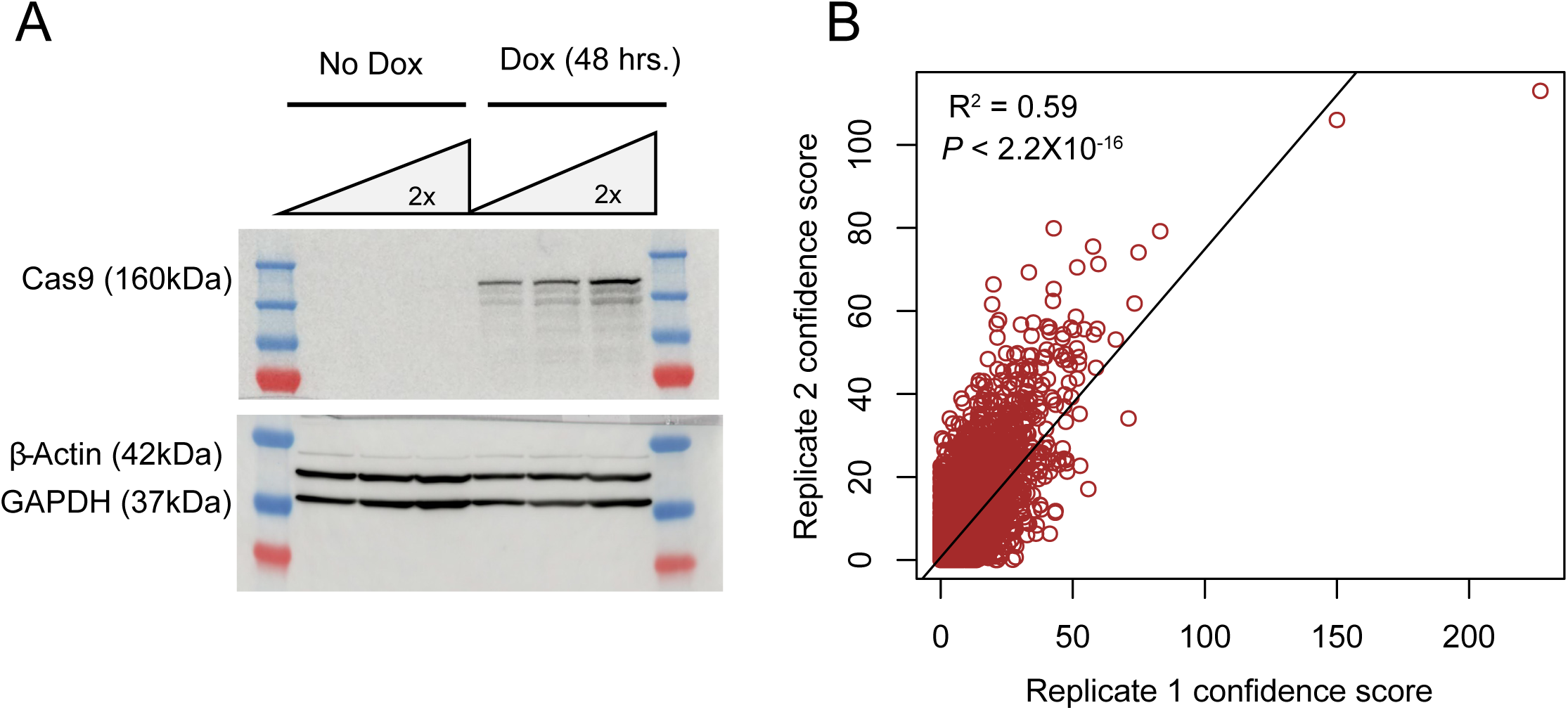
Cell line validation and replication of pigmentation CRISPR-Cas9 screen. (**A**) Western blot for Cas9-MNT1 cells showing Cas9 expression upon 48-hour doxycycline treatment. (**B**) Reproducibility of two genome-wide independent screens (n=2). CasTLE analysis of genome-wide screens in MNT1 cells, with CasTLE confidence score of 20,488 genes represented as individual points, analyzed by CasTLE likelihood ratio test. R^2^ and *P* value determined by linear regression.

**Fig. S3.**
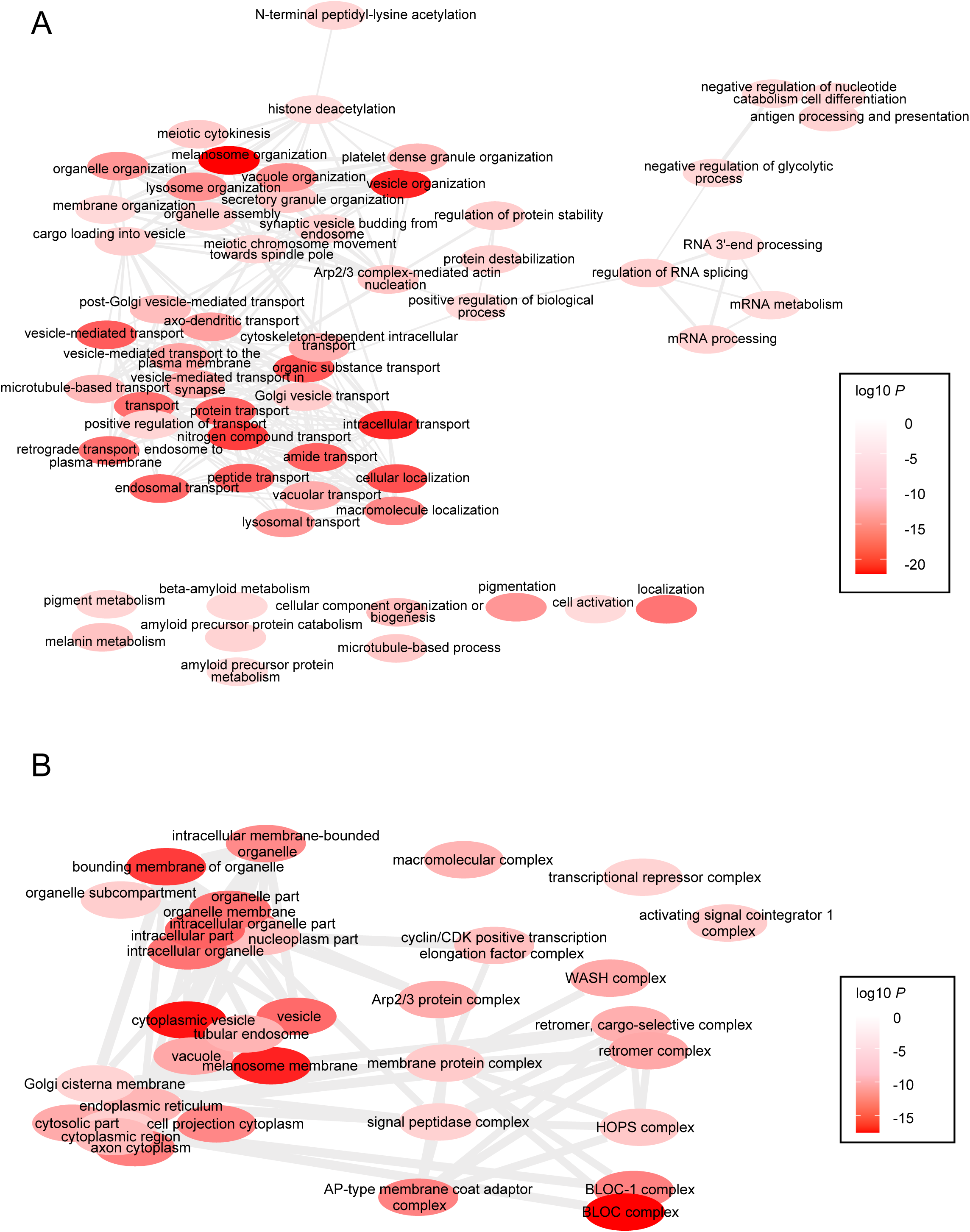
Functional classification of pigmentation screen hits. Gene ontology enrichment analysis for (**A**) biological processes and (**B**) biological components performed using GOrilla (*32*). The screen hits at 10% FDR were used as input gene set against all genes as background set. Enriched GO terms with q value <0.05 were used as an input in REVIGO (*33*) to remove redundant GO terms and make graphs. Similar GO terms are joined by the edges in the graph, where the line thickness indicates the degree of similarity. Color shading of bubbles represents hypergeometric test *P* values from GOrilla output.

**Fig. S4.**
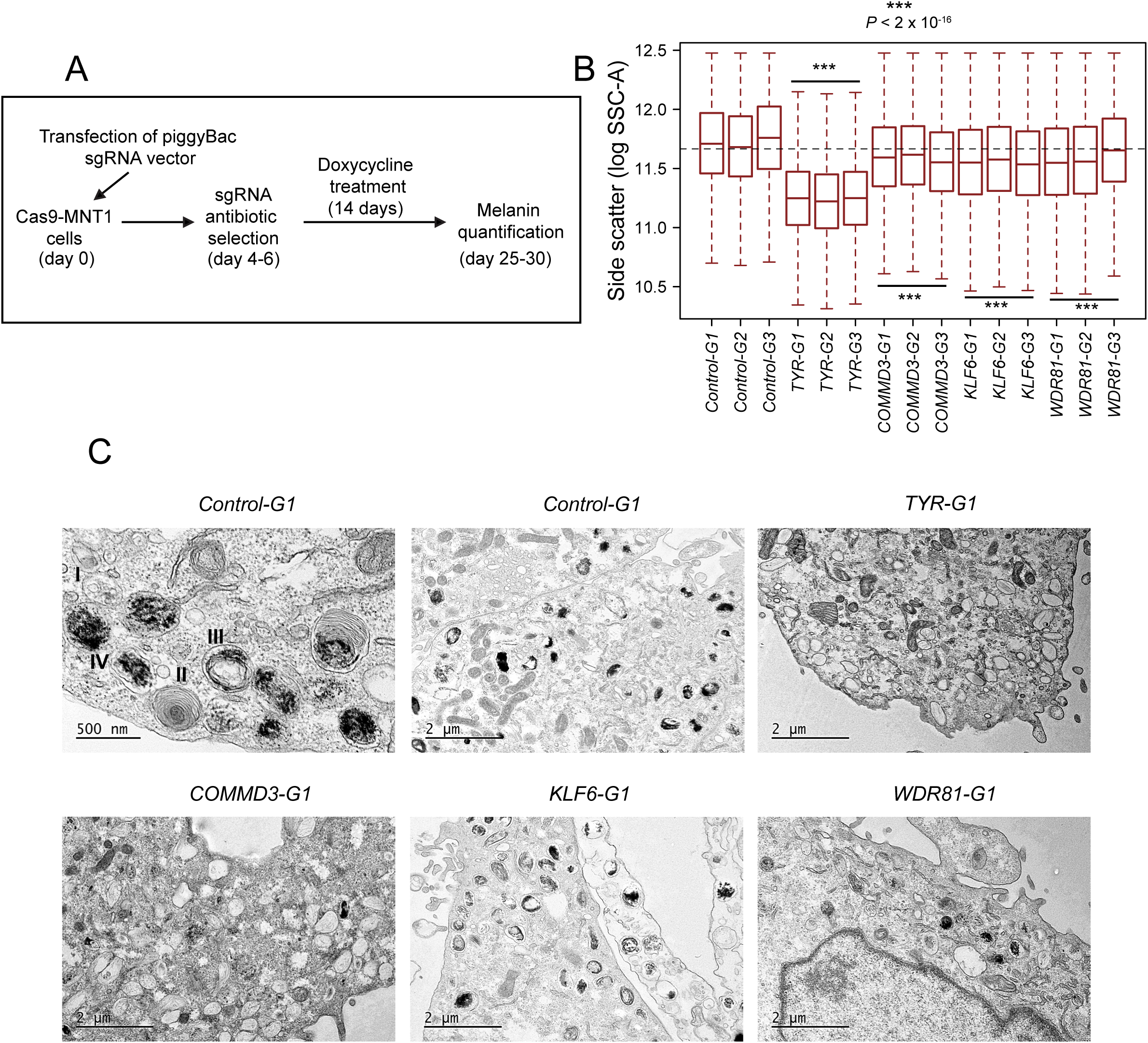
Effects of candidate gene knockouts on SSC and melanosome biogenesis. (**A)** Schematic of the CRISPR-Cas9 based validation of novel pigmentation genes. (**B)** Knockouts of candidate genes affect side scatter properties of the cells. Box plots show median log SSC and IQR, whiskers are 1.5 × IQR. Two-sided pairwise Wilcoxon Rank Sum Test with Benjamini & Hochberg correction was performed to compare SSC distributions and significance. *P* values shown are relative to control-edited cells. **(C)** TEM images of indicated gene knockout cells showing distribution of melanosomes.

**Fig. S5.**
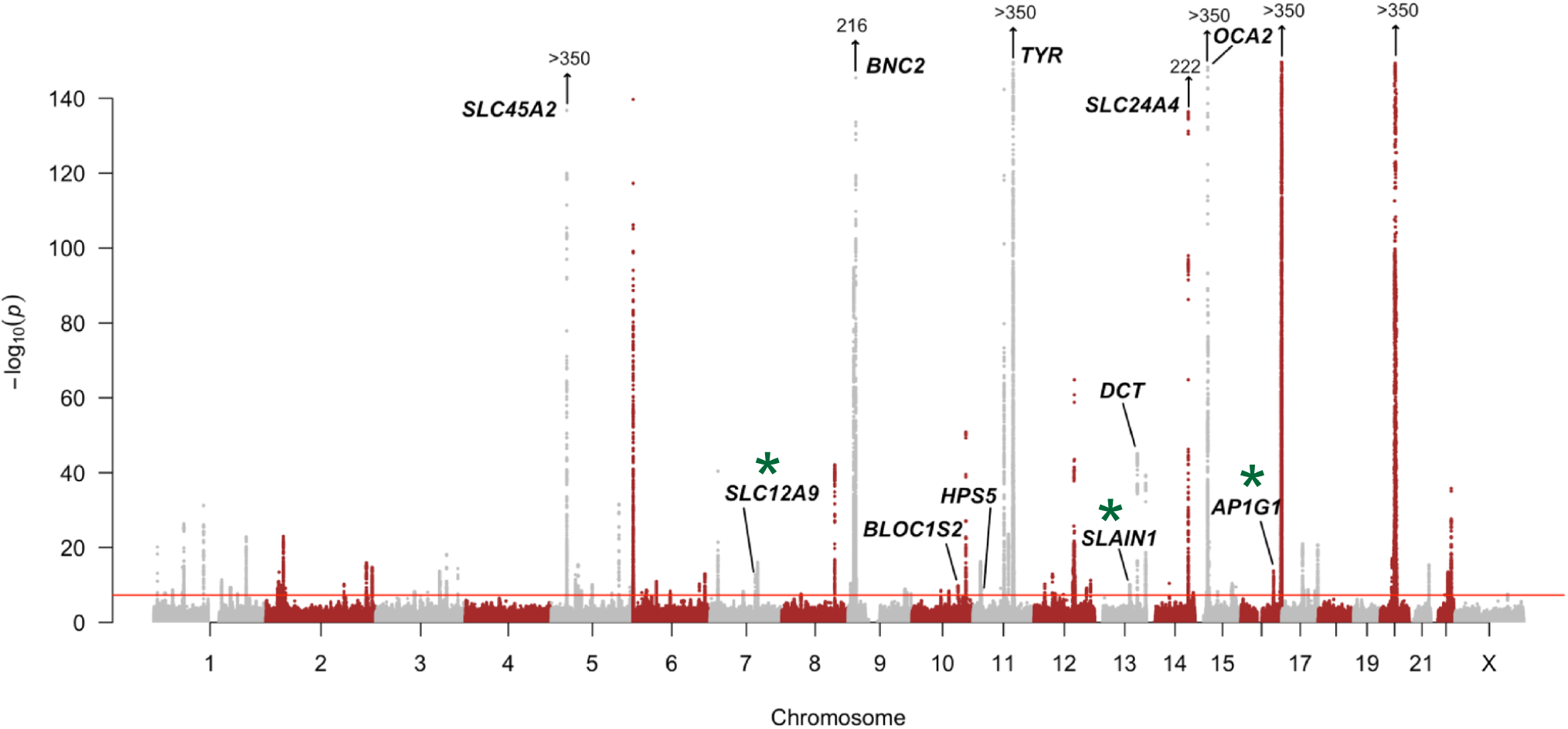
Select screen hits overlap skin color GWAS loci. Manhattan plot of GWAS for skin color in the UK Biobank (UKBB). Genes in bold were detected as pigmentation screen hits and lie within 100kb of a genome-wide significant GWAS signal (*P* < 5×10^−8^) for skin color. Genes with an asterisk have no previously described role in pigmentation. Vertical arrows indicate additional SNPs with the indicated -log10(*P*) of association.

**Fig. S6.**
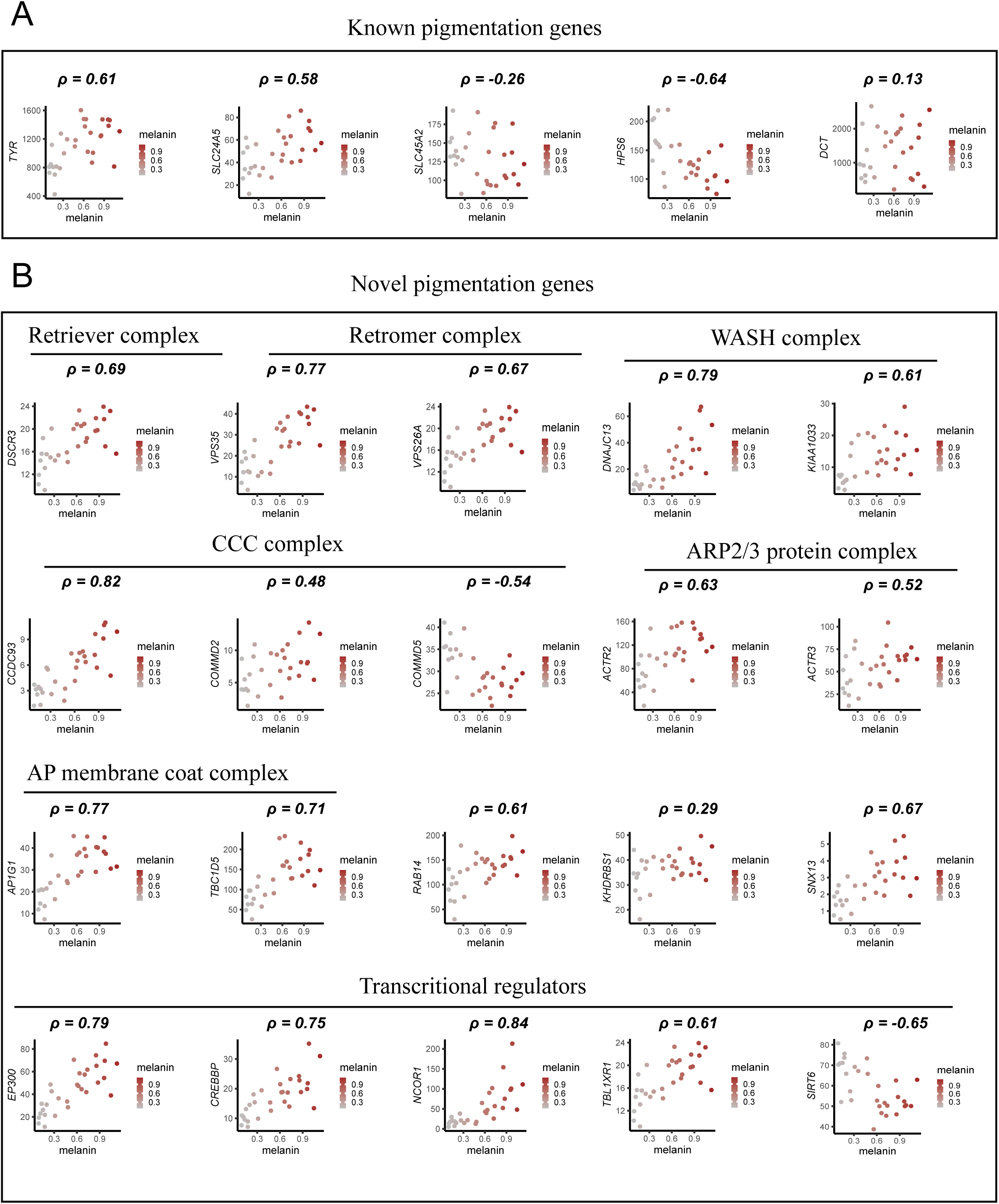
Spearman’s rank correlation coefficients (ρ) depicting relationship between melanin content and gene expression (TPM). (**A)** Known pigmentation genes rediscovered in the screen. (**B)** Novel melanin-promoting genes classified by their known molecular function. Plotted is melanin content (ordinate) vs RNA-seq expression level (TPM, abscissa).

**Fig. S7.**
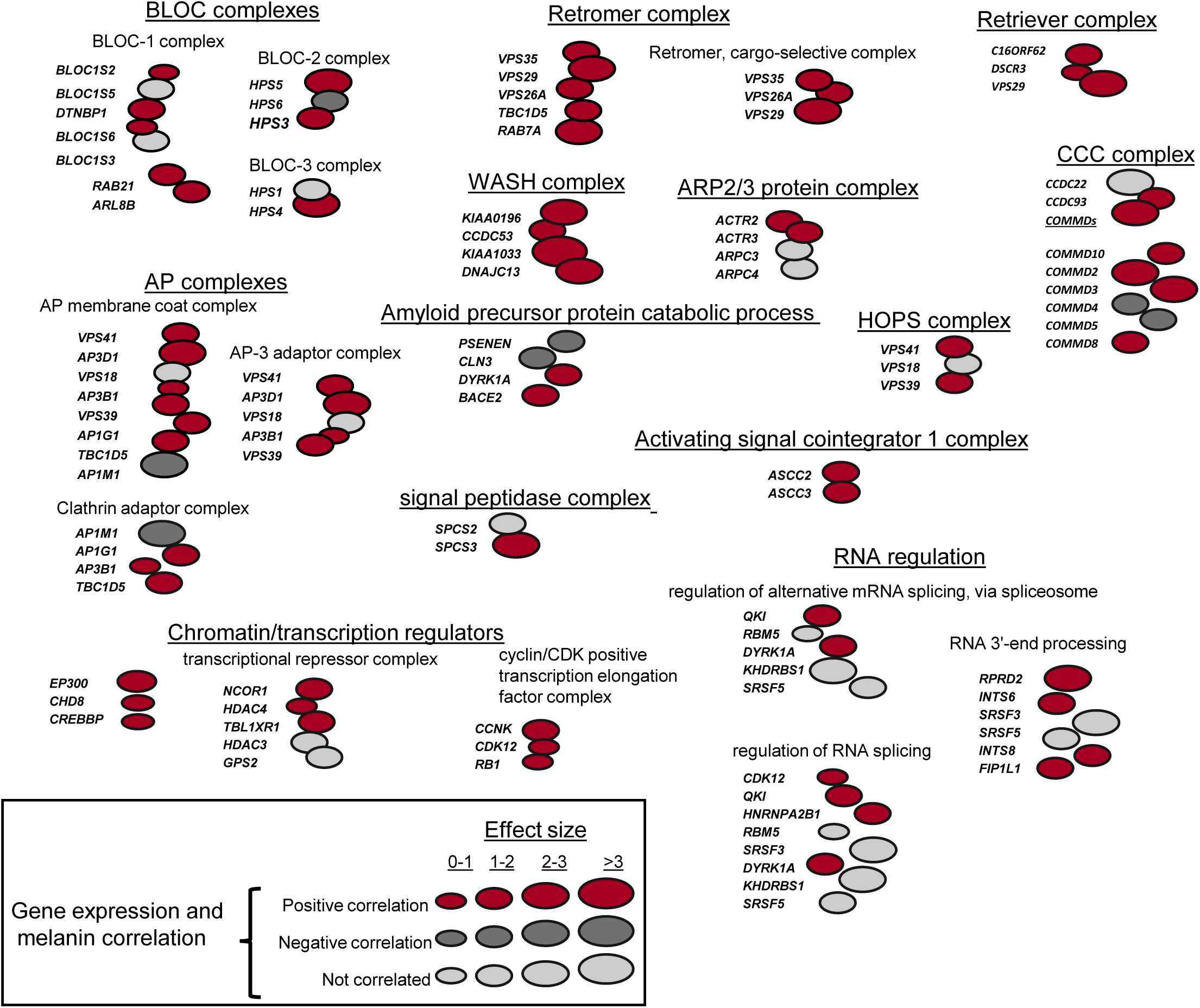
Classification of screen hits in relation to changes in gene expression in diversely pigmented MC. Schematic of screen hits grouped according to known biological function and presence in common macromolecular complexes (similar to Fig. 2E) is further color-coded based on the positive (dark red), negative (dark grey) or no (light grey) correlation between gene expression (TPM) and melanin content, as detected in the RNA-seq analysis of diversely pigmented individuals. Bubble size indicates CasTLE effect size.

**Table S1.** CRISPR screen hits corresponding to low side scatter FACS sort are shown with hits at 10% FDR are highlighted in yellow color and hits between 10-20% FDR are highlighted with blue color.

< Excel file>

**Table S2.** GO biological processes and components enrichment analysis for CRISPR screen hits at 10% FDR.

< Excel file>

**Table S3.** Fisher test contingency table and gene assignment table for human pigmentation heritability enrichment analyses.

< Excel file>

**Table S4.** Donor records for human foreskin tissue collection. Self-reported ancestry and melanin measurements are documented.

< Excel file>

**Table S5.** RNA-seq transcript per million (TPM) data for all the primary human melanocytes. All columns contain donor identifier as described in table S4.

< Excel file>

**Table S6.** Spearman correlation coefficients calculation for RNA-seq TPM and melanin (OD 400 nM) measurements.

< Excel file>

